# A neurometabolic mechanism involving dmPFC/dACC lactate in physical effort-based decision-making

**DOI:** 10.1101/2024.05.02.592220

**Authors:** N. Clairis, A. Barakat, Jules Brochard, Lijing Xin, C. Sandi

**Affiliations:** Laboratory of Behavioral Genetics, Brain Mind Institute, École Polytechnique Fédérale de Lausanne (EPFL), Lausanne, Switzerland; Transdisciplinary Research Areas, Life and Health, University of Bonn, Bonn, Germany; Center for Biomedical Imaging (CIBM), École Polytechnique Fédérale de Lausanne (EPFL), Lausanne, Switzerland; Institute of Physics (IPHYS), École Polytechnique Fédérale de Lausanne (EPFL), Lausanne, Switzerland

**Keywords:** Effort, physical effort, decision-making, motivation, ^1^H-MRS, fMRI, plasma, lactate, anterior insula, dorsomedial prefrontal cortex, dorsal anterior cingulate cortex, dmPFC, dACC

## Abstract

Motivation levels vary across individuals, yet the underlying mechanisms driving these differences remain elusive. The dorsomedial prefrontal cortex/dorsal anterior cingulate cortex (dmPFC/dACC) and the anterior insula (aIns) play crucial roles in effort-based decision-making. Here, we investigate the influence of lactate, a key metabolite involved in energy metabolism and signaling, on decisions involving both physical and mental effort, as well as its effects on neural activation. Using proton magnetic resonance spectroscopy and functional MRI in 63 participants, we find that higher lactate levels in the dmPFC/dACC are associated with reduced motivation for physical effort, a relationship mediated by neural activity within this region. Additionally, plasma and dmPFC/dACC lactate levels correlate, suggesting a systemic influence on brain metabolism. Supported by path analysis, our results highlight lactate’s role as a modulator of dmPFC/dACC activity, hinting at a neurometabolic mechanism that integrates both peripheral and central metabolic states with brain function in effort-based decision-making.

## Introduction

The ability to choose and execute effortful actions varies widely among individuals, reflecting differences in motivation levels and impacting health outcomes [1–3]. Understanding the neurobiological substrates underlying such variations is crucial, particularly as this knowledge can help address motivational dysregulation observed in neuropsychiatric and neurological disorders such as depression and Parkinson’s disease [4–6]. The dorsomedial prefrontal cortex/dorsal anterior cingulate cortex (dmPFC/dACC) and the anterior insula (aIns) play pivotal roles in guiding decisions that require effort [7, 8]. Despite their importance, the neurobiological mechanisms that govern their differential recruitment and how these mechanisms influence individual differences in motivational processes remain unknown.

Emerging evidence underscores brain metabolism as a crucial regulator of neural and cognitive functions [9–11], including motivated behaviors [12–14]. While glucose has been traditionally considered the primary energy source for brain function, including demanding cognitive processes [15], recent studies reveal that neurons also depend on alternative substrates, such as lactate, to support neural function and meet cognitive demands [16–19]. Lactate is a crucial brain metabolite which is primarily produced in astrocytes [20, 21] and muscle tissue during physical exercise [22–24] from where it may reach the brain through blood circulation [25, 26]. Initially regarded as a mere metabolic byproduct requiring clearance [27, 28] and associated with physical fatigue [29, 30], lactate’s ability to cross the blood brain barrier [31, 32] uniquely positions it to influence brain function and cognition [33, 34]. This dual origin and the transfer of lactate from peripheral sources to the brain underscore its potential to impact decision-making processes, particularly those involving effort. Beyond its metabolic functions, lactate acts as a signaling molecule [35, 36], influencing neuronal survival, plasticity, and memory formation [37–39]. Conversely, elevated lactate levels may disrupt cellular and metabolic balance, requiring regulation to maintain neuronal efficiency and prevent dysfunction [11, 40–43].

The brain must have evolved the ability to sense and interpret systemic energy levels – particularly signals reflecting the plentifulness or scarcity of metabolic energy resources required to support costly effortful behaviors, which could subsequently be incorporated in the evaluation of which actions to initiate during effort-based decision-making. Lactate’s potential as a signal of peripheral fatigue and its unique permeability across the blood-brain barrier suggest it is well-positioned to influence these neural computations. Given the emerging role of lactate as both a metabolic substrate and a potential neuromodulator, exploring its influence on the neurobiological mechanisms underpinning effort-based decision-making is essential. A crucial prediction is that individual differences in baseline lactate levels within specific brain regions—particularly dmPFC/dACC and aIns—will influence effort-based decision-making processes. Thus, we hypothesize that individual variation in lactate levels in the dmPFC/dACC and/or aIns integrate information about energy availability, regulating neural activity, thereby affecting effort-based decision-making. Such findings would provide a neurochemical basis to our computational understanding of effort-based decision making, offering a mechanistic explanation for interindividual variability in these behaviors.

To address these gaps and test our hypotheses about lactate’s role, we conducted a study using proton magnetic resonance spectroscopy (^1^H-MRS) in a 7 Tesla scanner and measured lactate levels in both the dmPFC/dACC and the aIns of 75 human participants. Functional magnetic resonance imaging (fMRI) data were then acquired as subjects engaged in a decision-making task, allowing quantification of brain activation patterns during effort-based decision-making. We found that dmPFC/dACC lactate levels are closely linked to neural activity during decision-making processes and, consequently, to individual behavioral choices. We also used computational modeling to extract key behavioral parameters driving motivated behavior and show that dmPFC/dACC lactate’s impact on neural activity and behavior was driven by its influence on the individual sensitivity to physical effort. While significant findings were observed in the dmPFC/dACC, they were not replicated in the aIns. Further exploration extended to examining plasma lactate concentrations, revealing their association with dmPFC/dACC lactate levels. Critically, path mediation analyses indicated that the relationship between plasma and dmPFC/dACC lactate levels and physical effort-based decision-making was mediated by dmPFC/dACC neural activity at the inter-individual level. This dual central-peripheral neurometabolic approach advances our understanding of lactate’s role in neural mechanisms underlying effort-based decision-making and identifies the dmPFC/dACC’s as a potential locus of control linking metabolic states and cognitive functions in humans.

## Results

### Behavioral Results

Participants engaged in a choice task structured into four blocks of 54 effort-based decision-making trials each, conducted concurrently with fMRI data acquisition. Each trial presented two options, each associated with different levels of effort and monetary incentives (**Fig. 1**). Each block focused on a specific type of effort: physical (squeezing a handgrip at 55% of their maximal force for varying durations) or mental (completing a set of correct responses in a 2-back task within a designated time), while monetary incentives were potential rewards or losses interleaved across trials. Both gains and losses were included in the task to account for the fact that individuals may display a ‘loss aversion’ bias [44], which would predict that they are ready to invest more effort for monetary losses than for equivalent monetary gains and the fact that both dmPFC/dACC and aIns have been regularly associated to processing aversive stimuli which could imply that they correlate with punishment sensitivity rather than effort [45, 46]. During the choice, one option was consistently associated with a fixed low-effort difficulty and low monetary incentive (i.e., either a low reward or a high potential loss). The alternative option was always linked to a higher monetary incentive (i.e., either a higher reward or a smaller loss) paired with a high effort. The incentive and effort levels associated with the high-effort option varied independently across trials. On each trial, participants were required to select their preferred option based on their subjective preferences, along with their confidence (high or low) in their decision. After one option had been selected, participants were required to exert the effort chosen on each trial to secure the specified reward or to prevent the doubling of the indicated loss.

**Figure 1:**
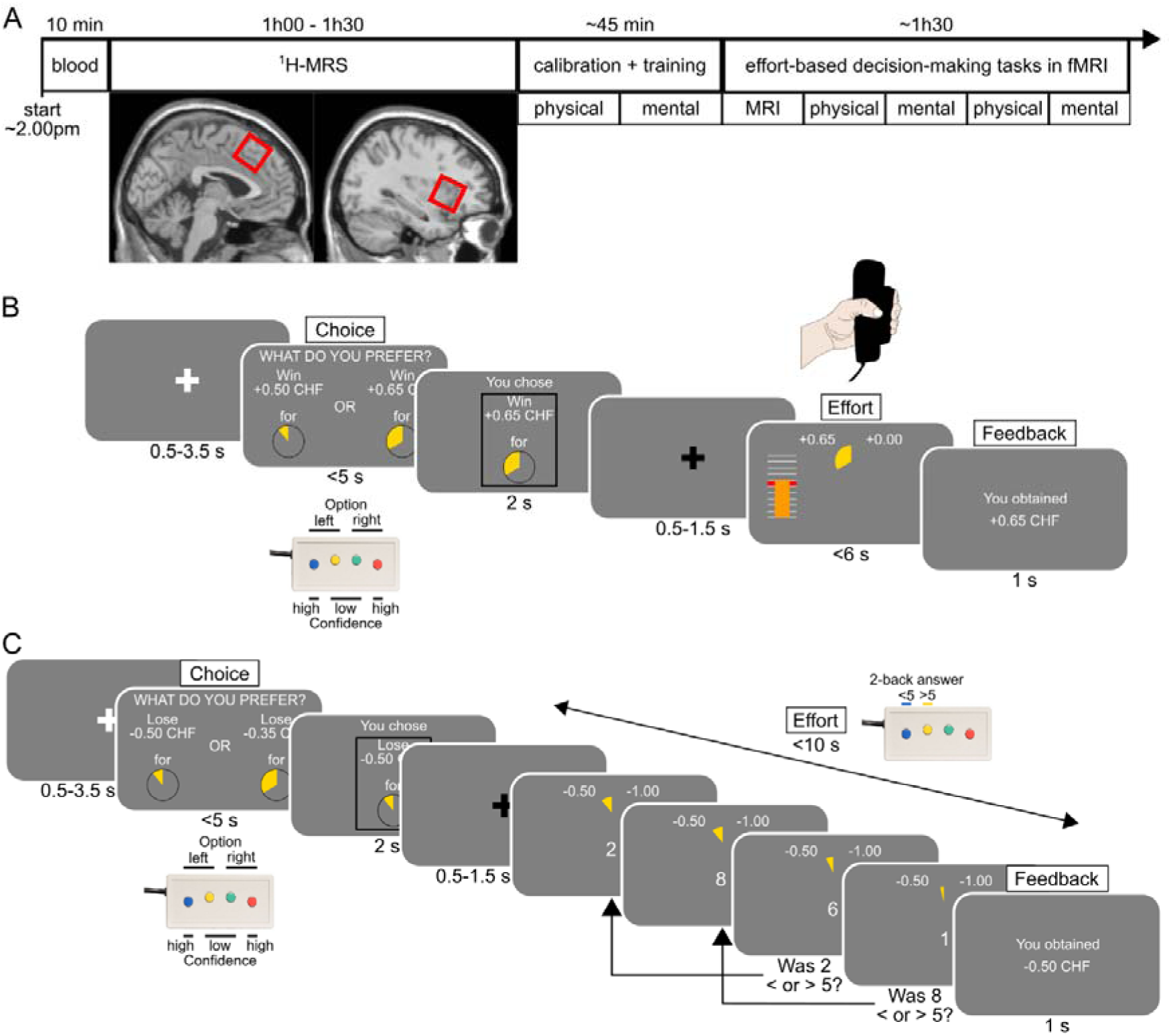
Behavioral task. **A]** Experimental timeline. Participants first started with a blood extraction in a dedicated medical facility followed by proton magnetic resonance spectroscopy (H-MRS) in a magnetic resonance imaging (MRI) scanner to measure lactate in the dorsomedial prefrontal cortex/dorsal anterior cingulate cortex (dmPFC/dACC) (left) and the anterior insula (aIns) (right). The intended voxel placement for each region of interest is highlighted by a red square, superimposed on the SPM single subject T1 template (see **Fig. S1-2A** for the actual average voxel location across subjects). This measure allowed us to acquire lactate, characterized by a doublet around ∼1.33ppm, in both dmPFC/dACC and aIns (**Fig. S1-2B**). Next, calibration and training outside the MRI scanner allowed us to adjust effort and monetary incentive levels, based on individual indifference points. Participants then returned in the MRI scanner to perform the task during fMRI in blocks of physical or mental effort (order counterbalanced), with blocks comprising 54 trials and lasting ∼5-15 minutes each. Before and after each block, participants were asked to perform their maximal performance again. **B-C]** Effort-based decision-making tasks. Subjects first performed a choice between the two options, always followed by the exertion of the effort selected in order to obtain the money selected (in the reward trials) or to avoid losing twice the amount selected (in the punishment trials). Reward and punishment trials were interleaved. The proportion of high effort choices and deliberation times depending on the incentive and on the effort levels can be found in **Fig. S1-1**. **B]** Physical effort task. The physical effort consisted in squeezing a handgrip with the left hand above 55% of the maximal voluntary contraction (MVC) force for variable durations based on the effort selected. **C]** Mental effort task. The mental effort consisted in completing a 2-back task within 10s. The 2-back task required to indicate if the number displayed two digits back was below or above the number five. The number of correct answers to provide during each trial depended on the maximal number of correct responses (MNCR) obtained during calibration and on the effort selected.

Despite calibration around each task’s indifference point (see *Supplementary Methods*), participants showed a stronger preference (p < 0.001) for the high effort (HE) option in the mental task (74.97 ± 2.04%) compared to the physical task (58.24 ± 2.33%) on average. Preference for the HE option in both tasks diminished with less appetitive monetary incentives or increased effort requirements (**Fig. S1-1A**). Surprisingly, instead of loss aversion, our participants tended to value gains more than losses in our tasks (see *Supplementary Results*). The task design was optimized to assess individual aversion to effort rather than the subjective estimation of success, i.e. risk [47], which is often confounded with effort [48]. This was done by calibrating each task to the individual’s maximal capacity and setting difficulty levels allowing participants to maintain high performance consistently throughout the task (see *Supplementary Methods*). Indeed, participants sustained high average performance levels in both tasks (98.26 ± 0.36% for mental efforts and 98.64 ± 0.45% for physical efforts), confirming their ability to meet the task demands on most trials. This high performance effectively rules out risk discounting as a confounding factor both within and between the tasks.

Participant choices were analyzed using a computational model that incorporated both external factors (monetary incentive, effort level) and internal states (physical fatigue, momentary mental efficiency). Simulations confirmed parameter recoverability and identifiability (see *Supplementary Results*). Furthermore, a conventional generalized linear model (GLM) validated the significant impact of each model variable on choice behavior and deliberation times (see *Supplementary Results*), corroborating that both monetary incentives and effort difficulty influenced decision-making and deliberation times in our task (**Fig. S1-1**).

### Brain and Plasma Lactate Levels in Relation to Effort-Based Decision-Making

After validating our experimental paradigm, we examined the relationships between lactate levels in the dmPFC/dACC and the aIns (**Fig. S1-2A**), respectively, and decision-making variables in the behavioral tasks. Resting lactate levels in the dmPFC/dACC, measured using H-MRS prior to task initiation, were negatively correlated with the propensity to choose the HE option across tasks (**Fig. 2A**; r = -0.322, p = 0.013), indicating that higher lactate concentrations in this region were associated with a reduced likelihood of selecting HE options. Conversely, lactate levels in the aIns did not show a significant relationship with HE choice frequencies (**Fig. 2A** ; r = -0.144, p = 0.358), suggesting regional specificity in lactate’s association with decision-making. Although there was a tendency for dmPFC/dACC and aIns lactate levels to correlate (**Fig. 2B** ; r = 0.236, p = 0.132), it did not reach statistical significance, further supporting the notion of regional variations in brain lactate levels [49]. However, when comparing the two correlations with a one-tailed Steiger’s test restricted to the 41 individuals who were not outliers in any of the three measures considered, the correlation coefficients were not significantly different (z = -0.879; p = 0.190), preventing us from drawing any conclusion on the specificity of our results. In summary, although our results suggest a strong role of the dmPFC/dACC in integrating energy-related signals into decision-making processes, based on our data, we cannot exclude that the aIns also plays a role in these processes when looking across both tasks.

**Figure 2:**
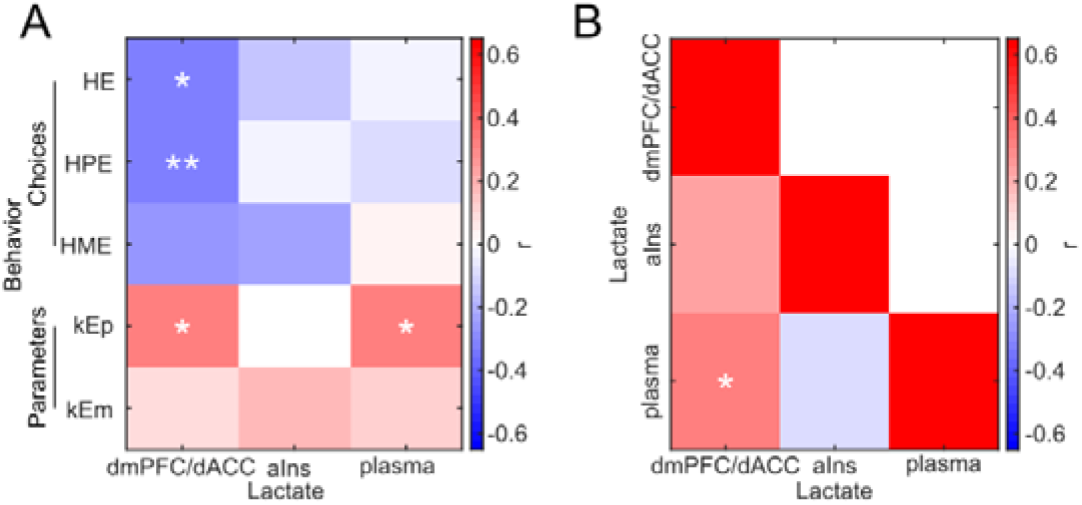
Correlations between plasma, dmPFC/dACC and aIns lactate levels and behavioral measures of motivation. **A**] Heatmap displaying Pearson correlation coefficients between dmPFC/dACC, aIns and plasma lactate levels and behavioral measures of motivation (high effort [HE], high physical effort [HPE] and high mental effort [HME] choices, and physical [kEp] and mental [kEm] effort sensitivities). Significance levels: ***p<0.001; **p<0.01; *p<0.05. **B**] Heatmap displaying Pearson correlation coefficients among lactate measurements across the dmPFC/dACC, the aIns, and plasma. Significance levels in both panels: ***p<0.001; **p<0.01; *p<0.05. We also verified that plasma and brain measures of lactate were not correlating with measures of capacity like maximum voluntary contraction force (**Fig. S2-1A**), the maximum number of correct responses in the mental task (**Fig. S2-1B**), subjective fatigue (**Fig. S2-1C**), or maximal performance (**Fig. S2-1D**). We also verified that there were no sex differences in behavior (**Fig. S2-2A**) or in plasma or brain levels of lactate (**Fig. S2-2B**). Finally, we also verified that plasma and brain levels of lactate were not correlated with sleep ratings (**Fig. S2-3**), with stress- and anxiety-related questionnaires in general and on the day of the experiment and with stress-related physiological variables such as cortisol (**Fig. S2-4**).

Regarding plasma lactate, there was no direct correlation between systemic lactate levels and HE choices (**Fig. 2A** ; r = -0.016, p = 0.906). However, a small yet significant positive correlation was observed between plasma lactate and dmPFC/dACC lactate levels (**Fig. 2B**; r = 0.314, p = 0.015). This correlation did not hold for the aIns (**Fig. 2B** ; r = -0.091, p = 0.563), which had additionally a significantly lower correlation with plasma lactate levels than the one for the dmPFC/dACC (note, again, that this comparison is based on a one-tailed Steiger’s test restricted to the 43 individuals present in all measures; z = 1.887, p = 0.030), indicating a unique connection between systemic and dmPFC/dACC lactate levels, but not with the aIns.

Analysis of task-specific choices revealed differential correlations across the two brain regions. Specifically, dmPFC/dACC lactate levels were negatively correlated with the selection of high physical effort (HPE) options (**Fig. 2A** ; r = -0.332, p = 0.010) and showed a non-significant trend for high mental effort (HME) choices (**Fig. 2A** ; r = -0.249, p = 0.058). In contrast, aIns lactate levels did not significantly correlate with either HPE (**Fig. 2A**; r = -0.020, p = 0.897) or HME (**Fig. 2B**; r = -0.225, p = 0.147) choices. Moreover, the dmPFC/dACC-lactate/HPE choices correlation coefficient was significantly more negative than the aIns lactate/HPE correlation coefficient (note, again, that this comparison is based on a Steiger’s test restricted to the 42 individuals with no outliers in any of these measures; z = -1.887, p = 0.029) confirming dmPFC/dACC’s specific role in integrating energy-related signals into physical effort-based decision-making. Interestingly, the comparison of the correlation coefficients of the dmPFC/dACC-lactate/HPE choices correlation and the dmPFC/dACC-lactate/HME choices correlation showed that they were not significantly different (z = -0.505, p = 0.307) which further suggests that dmPFC/dACC lactate levels may also play a role in mental effort decisions, although this role may be more subtle than for physical efforts. In summary, individuals with higher dmPFC/dACC, but not aIns, lactate levels tend to avoid energetically costly physical efforts, suggesting that lactate may modulate the threshold for engaging in high-effort tasks by influencing neural circuits responsible for effort evaluation.

Our computational model allowed us to reveal that these effects might be related to differences in the individual sensitivities for physical effort (kEp). Indeed, a positive correlation was found between dmPFC/dACC lactate levels and kEp (**Fig. 2A**; r = 0.317, p = 0.017), indicating that individuals with higher lactate levels in this region display an increased sensitivity to physical effort. This correlation seemed specific to kEp, as no significant correlations were observed with other behavioral parameters such as the equivalent effort sensitivity in the mental domain (kEm; **Fig. 2A**; r = 0.077, p = 0.560), the sensitivity to physical fatigue (kFp; r = -0.008, p = 0.952), nor any of the other behavioral parameters (all p > 0.4). However, note that when performing 2-by-2 Steiger tests between the correlation coefficients, except for the correlation between dmPFC/dACC-lactate and the kBias parameter which was significantly less robust than the dmPFC/dACC-lactate/kEp correlation (z = 2.632, p = 0.004), the difference between the correlation coefficient of the dmPFC/dACC-lactate/kEp correlation and the correlation coefficient for the other parameters [e.g., dmPFC/dACC-lactate/kEm correlations (z = 1.194, p = 0.116) or dmPFC/dACC-lactate/kFp correlation (z = 1.303, p = 0.096)] was not significant. Although the dmPFC/dACC-lactate/kEp correlations always tended to be stronger than the respective dmPFC/dACC-lactate correlations with the other parameters (all z ∈ [1.1 1.6], all p ∈ [0.06 0.13]), this prevents us from drawing strong conclusions regarding the specificity of our correlations with kEp. It also suggests that dmPFC/dACC lactate levels may also play a role in mental effort decisions, although this role may be more subtle than for physical efforts. There were also no significant correlations between aIns lactate levels and kEp (**Fig. 2A**; r = 0.007, p = 0.963), nor any of the other parameters (all p > 0.08). However, the direct comparison of dmPFC/dACC-lactate/kEp to the aIns-lactate/kEp correlation coefficients was not significant when restricting the comparison to the individuals present in both groups (z = 1.267, p = 0.103), preventing us from drawing any strong inference on the dmPFC/dACC specificity. Interestingly though, plasma lactate levels correlated positively with kEp (**Fig. 2A** ; r = 0.320, p = 0.014) and with dmPFC/dACC lactate levels (**Fig. 2B** ; r = 0.314, p = 0.015) suggesting that plasma lactate may impact the sensitivity to physical effort via its influence on dmPFC/dACC lactate levels. Notably, plasma lactate did not correlate with any of the other behavioral parameters of our model (all p > 0.1) further confirming that dmPFC/dACC and plasma lactate only correlate significantly with the sensitivity to physical effort. We also performed a few additional controls regarding the potential link between plasma and brain levels of lactate and physical capacity (**Fig. S2-1**), sex differences (**Fig. S2-2**), sleep (**Fig. S2-3**) and stress- and anxiety-related variables (**Fig. S2-4**) which all confirmed that lactate was solely correlated with differences in physical effort motivation but not in physical capacity, sex differences, sleep or stress (see *Supplementary Results*). Notably, although we did not observe any correlation with fatigue-related behavioral components and lactate in our task, our results do not preclude the possibility that exercise-driven lactate increase could also impact central fatigue since we did not monitor lactate levels changes during the task. Brain regions such as the dmPFC/dACC, responsible for deciding whether to engage and persevere in effortful actions [50, 51] and which activity varies with fatigue [52], may monitor elevated lactate levels as part of their computation in determining whether to engage in such actions. Interestingly, baseline resting-state brain levels of lactate appear to be strikingly stable across subsequent sessions [53] or between morning and afternoon [54], suggesting that the results we observe really correspond to trait differences in lactate and effort sensitivity, and not just to temporary state levels of lactate. In other words, the global body metabolism efficiency, reflected in peripheral and dmPFC/dACC lactate levels, may drive a general trait sensitivity to physical efforts that we capture in our experiment and that is not just due to momentary fluctuations in lactate levels.

### Neural Activity and Effort-Based Decision-Making

After establishing that dmPFC/dACC lactate levels, but not aIns, correlate with physical effort motivation, we further explored if dmPFC/dACC neural activity during decision-making correlates with intra- and inter-individual differences in physical effort-based decision-making. We assessed dmPFC/dACC activity at the time of choice, hypothesizing it to represent an energization signal [50, 55] modulating the proportion of HE choices based on the effort required. Alternative accounts of the dmPFC/dACC during decision-making relate it to negative subjective value encoding [46, 56–58] and to deliberation times [59–61]. We therefore included these regressors in our analysis.

Using a generalized linear model (GLM1) with the level of chosen effort (Ech), the subjective value of the chosen option (SVch), and deliberation times (DT) as parametric modulators of the choice regressor, we observed significant dmPFC/dACC activity correlating with Ech when pooling the data across both physical and mental tasks (**Fig. 3A**). These findings underscore the dmPFC/dACC’s central role in supporting effort-based decision-making processes and corroborates its potential role in encoding an energization signal [50]. Additionally, a smaller cluster was identified in the left (but not right) aIns when looking at Ech correlates across physical and mental effort tasks with a slightly more lenient threshold (**Fig. 3A**), suggesting that the left aIns might also be involved in supporting effort-based decisions. Interestingly, the dmPFC/dACC also seemed to correlate negatively with SVch and positively with DT (**Fig. S3-2** & *Supplementary Results*) in agreement with the literature and the correlation held independent of the way the parametric modulators of choice (Ech, SVch, DT) were orthogonalized to each other (**Fig. S3-3** & Supplementary Results). Therefore, we validated the robustness of our tasks not only for assessing effort-based decision-making across physical and mental domains as shown above (*Results: Behavioral Results*), but also by confirming their ability to engage the dmPFC/dACC and aIns during such processes. Furthermore, we also controlled that dmPFC/dACC and aIns reflected the level of the chosen effort [50, 56], and therefore were potentially determining the choice to make, and not just the effort costs of the HE option (**Fig. S3-4** & *Supplementary Results*) as has been suggested [8, 62]. This supports the documented contributions of these regions to effort valuation and decision-making [7, 8, 56, 62–66], with our analysis distinguishing this activation from factors like subjective value perception [46, 57, 58] or reaction times [59–61].

**Figure 3:**
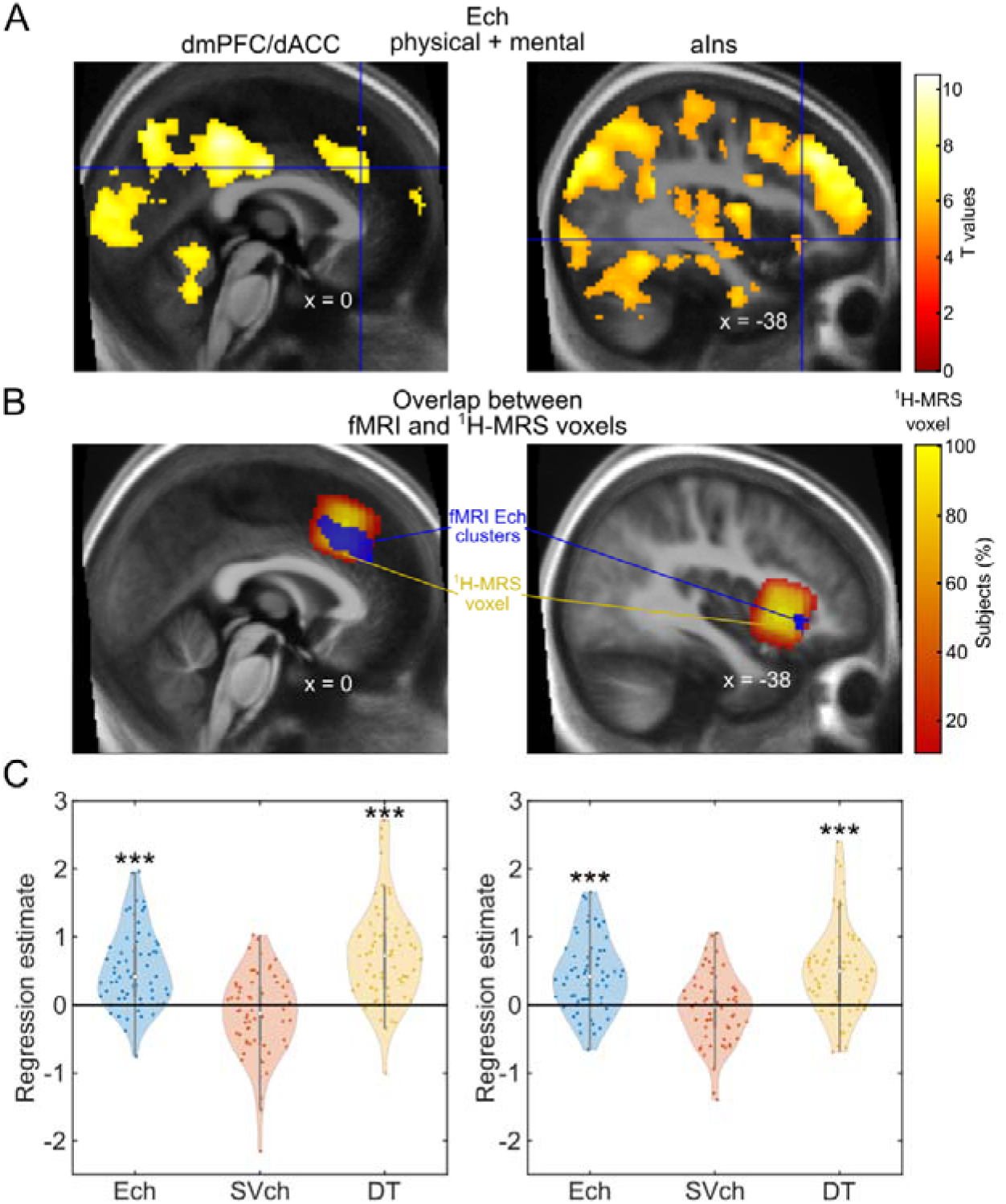
Task-Related Neural Activity and Spectroscopy Voxel Overlap. **A]** Brain activation related to the level of the chosen effort (Ech) across tasks. A t.test against zero across participants (N = 63), reveals a positive correlation between dmPFC/dACC (left), voxel-wise thresholded at p < 0.001 after family-wise error (FWE) correction for multiple comparisons (cluster peak: x = 0, y = 32, z = 32 in MNI coordinates; cluster size: k = 686 voxels; cluster p.value after family-wise error (FWE) correction for multiple comparisons: p < 0.001), and aIns (right), voxel-wise thresholded at p < 0.05 FWE-corrected (x = -38, y = 26, z = -4; k = 55; cluster p.value after FWE correction for multiple comparisons: p < 0.001), activities and Ech. No cluster correlating with Ech was found in the right aIns. The Neurovault full map can be found at: https://neurovault.org/collections/17029/. **B]** ^1^H-MRS voxels overlap with the fMRI clusters. The left panel highlights the overlap between the dmPFC/dACC fMRI cluster from [**A**] and the ^1^H-MRS dmPFC/dACC density map across subjects (details in **Fig. S1-2**). The overlap between the dmPFC/dACC fMRI cluster for Ech and the dmPFC/dACC ^1^H-MRS voxel location revealed that 99.6% of the voxels of the dmPFC/dACC fMRI cluster and 14.1% of the dmPFC/dACC ^1^H-MRS measurement were overlapping, confirming that our ^1^H-MRS voxel placement was highly precise and relevant for the purpose of this study. Even when restricting the analysis to the ^1^H-MRS voxels present in at least 90% of the participants, there was still an important overlap of 38.3% of the dmPFC/dACC voxels with 49.4% of the dmPFC/dACC H-MRS voxels, confirming the correct placement of our voxel of interest. The right panel highlights the overlap between the left aIns fMRI cluster from [**A**] and the ^1^H-MRS aIns density map across subjects (details in **Fig. S1-2**). The overlap entailed 100% of the fMRI cluster and 0.98% of the aIns H-MRS voxel when including all subjects, and 27.3% of the aIns fMRI cluster and 3.7% of the H-MRS voxel when restricting to the H-MRS voxels sampled in at least 90% of all the participants, further confirming that our ^1^H-MRS aIns placement was also meaningful for studying Ech. **C]** Regression estimates for effort chosen (Ech), the subjective value of the chosen option (SVch) and deliberation time (DT) during the choice period within the H-MRS dmPFC/dACC (N = 63, left panel) and the H-MRS aIns (N = 60, right panel). The dmPFC/dACC activity was positively associated to Ech (β = 0.533 ± 0.072; p < 0.001), a relationship holding even when splitting the data between physical and mental effort tasks (**Fig. S3-1**). The dmPFC/dACC activity was also positively associated to DT (β = 0.738 ± 0.090; p < 0.001) and it also tended to be negatively associated to SVch (β = -0.147 ± 0.076; p = 0.057) and. The overlap between the three dmPFC/dACC fMRI clusters can be observed in **Fig. S3-2**. The aIns activity was also positively associated to Ech (β = 0.441 ±0.071; p < 0.001) in both physical and mental tasks (**Fig. S3-1**), as well as with DT (β = 0.500 ± 0.084; p < 0.001), but not with SVch (β = -0.052± 0.067; p = 0.437). Note that the ROI results are stable even if regressors are orthogonalized to each other (**Fig. S3-3**). Furthermore, the observed outcome is really tied to an increase with the level of the effort chosen and not with a computation of the effort costs related to the high effort option (**Fig. S3-4**). Significance of Student t-tests against zero: ***p<0.001; **p<0.01; *p<0.05.

Additionally, an overlap analysis confirmed a significant correspondence between the dmPFC/dACC fMRI cluster related to Ech and the dmPFC/dACC voxels measured by ^1^H-MRS (**Fig. 3B**), underscoring the accurate alignment of our imaging techniques with the neural processes involved in effort-based decisions. This was further confirmed by looking at the correlation between the activity of the ^1^H-MRS dmPFC/dACC and aIns voxels (hereafter called dmPFC/dACC and aIns) and Ech (**Fig. 3C**), which was also still present when looking at each task independently (**Fig. S3-1**).

Having confirmed the correlation between dmPFC/dACC and Ech at the intra-individual level, we also explored how differences in dmPFC/dACC Ech regression estimates could reflect inter-individual variations in physical effort decisions. Initially, we hypothesized that higher Ech dmPFC/dACC estimates would lead to increased energization and to a reduced kEp, therefore resulting in higher HPE choices. Alternatively, we considered that higher Ech estimates might indicate that more substantial dmPFC/dACC activation is required to reach the same level of motivation, suggesting that effort costs are perceived as higher in those individuals, thus discouraging high effort choices. This latter hypothesis aligns with the concept that dmPFC/dACC activity represents effort costs [51, 65, 67], influencing decision-making by modulating perceived effort expenditure. Negative correlations were found between the Ech regression estimates and the overall proportion of HE choices across participants, with both dmPFC/dACC (r = -0.335, p = 0.008) and aIns (r = -0.402; p < 0.001), indicating that higher neural sensitivity to effort choices is associated with a lower propensity to choose HE options (**Fig. 4**). This result was specific to the Ech regression estimate, as SVch and DT regression estimates in dmPFC/dACC and in aIns were not significantly correlated to the proportion of HE choices (**Fig. S4-1C-D** and *Supplementary Results*). Analyzing tasks independently, the relationship between the fMRI Ech regression estimate and motivated behavior held significant for HPE choices (**Fig. 4A-B**; dmPFC/dACC: r = -0.467, p < 0.001; aIns: r = -0.491; p < 0.001), but not for HME choices (**Fig. 4A**, **Fig. S4-1A**; dmPFC/dACC: r = -0.036, p = 0.780; aIns: r = -0.141; p = 0.286), highlighting task-specific differences in how effort cost influences decision-making.

**Figure 4:**
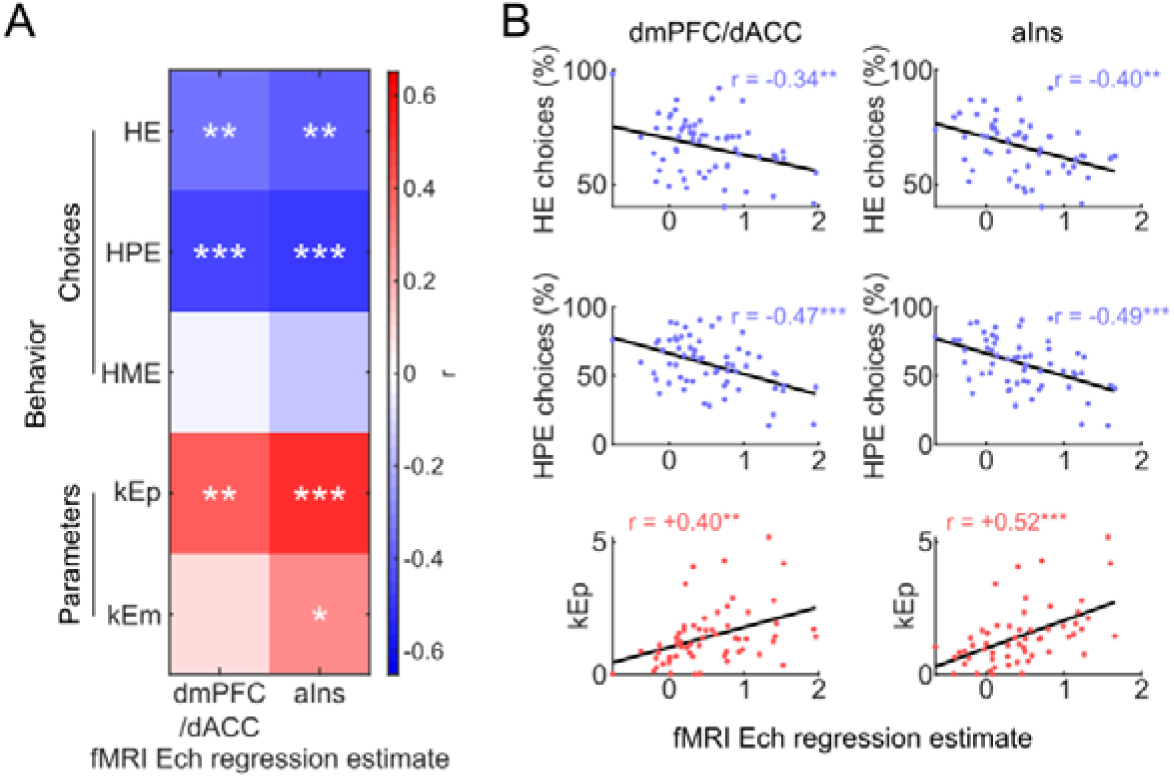
Correlation between behavioral motivation measures and dmPFC/dACC and aIns neural activity during choice. **A**] Heatmap displaying Pearson correlation coefficients between motivational behavioral variables (high effort [HE], high-physical effort [HPE], high mental effort [HME] and effort sensitivity parameters in the physical [kEp] and the mental [kEm] domain) against dmPFC/dACC and aIns effort chosen (Ech) regression estimate during decision-making. Significance levels: ***p<0.001; **p<0.01; *p<0.05. **B**] Scatter plots depicting the significant correlations from [**A**]. The plots highlight the linear relationship between dmPFC/dACC and aIns BOLD regression estimates for Ech across participants and behavioral measures of motivation (HE and HPE choices and kEp). Each point represents an individual participant. The scatter plots for the correlations with HME and kEm are displayed in **Fig. S4-1A** . The correlation heatmap for the other parameters from the model (kR, kP, kFp, kLm, kBias) is displayed in **Fig. S4-1B** . We also verified that none of the correlations displayed here was significant when replacing the Ech regression estimate by the SVch (**Fig. S4-1C**) or the DT (**Fig. S4-1D**) regression estimates.

In examining the specific contributions of dmPFC/dACC and aIns activities to decision-making processes involving physical effort rather than being driven by other task parameters, our analyses identified a clear link between neural activity in these regions and sensitivity to physical effort kEp (**Fig. 4**; dmPFC/dACC: r = 0.403; p = 0.001; aIns: r = 0.519; p < 0.001). Activity in the aIns (r = 0.277; p = 0.034), but not in the dmPFC/dACC (r = 0.087, p = 0.503) also correlated with the sensitivity to mental effort kEm (**Fig. 4A**, **Fig. S4-1A**), but none of the other parameters included in our model correlated with the Ech regression estimate in the dmPFC/dACC (**Fig. S4-1B**, all p > 0.25), nor in the aIns (**Fig. S4-1B**, all p > 0.1), highlighting the specific influence of dmPFC/dACC and Ins activity on effort sensitivity, with a particular emphasis on physical effort, particularly for the dmPFC/dACC.

In summary, our fMRI analysis indicates that dmPFC/dACC activity is closely associated with the selection of physical effort levels both at the intra- and at the inter-individual level, emphasizing its crucial role in the neural processes of effort valuation and motivation. This region’s activity patterns reflect individual variations in physical effort motivation and elucidate the cognitive substrates that account for differences in the propensity to engage in physically demanding tasks.

### Path Analysis: Lactate It/ Neural Activity It/ Effort-Based Decision-Making

Given the associations between both dmPFC/dACC lactate levels and fMRI activity during decision-making with the propensity to choose HPE options, but not with HME choices, we next examined whether dmPFC/dACC neural sensitivity to Ech serves as a mediator in the impact of dmPFC/dACC lactate levels on these choices. A mediation analysis aims to test whether a variable M statistically mediates the direct relationship between two correlated variables X and Y (path c) through the indirect path going from XlzlM (path a) to MlzlY (path b). If the indirect path is significant (max(p(a), p(b)) < 0.05) and the strength of the direct path is reduced after taking into account the indirect path (c’<c), then one can consider that M mediates the effect of X on Y. A first mediation analysis revealed that dmPFC/dACC Ech estimate significantly mediates the relationship between baseline dmPFC/dACC lactate levels and the selection of HPE options (Fig. 5A; mediation p = 0.004). Notably, when this neural activity mediator was considered, the direct influence of dmPFC/dACC lactate on HPE choices (c = -0.332, p = 0.010) diminished and became non-significant (c’ = -0.184, p = 0.148), suggesting that lactate affects HPE choices primarily through its impact on neural activity, thereby providing a direct mechanistic link between metabolic state and decision-making processes. The same result could be observed for kEp (Fig. 5B; mediation p = 0.035) with the direct path from dmPFC/dACC lactate to kEp initially significant (c = 0.317, p = 0.016) becoming non-significant after considering the fMRI mediator (c’ = 0.198, p = 0.151), confirming that higher lactate levels may reduce the motivation for physical efforts by influencing neural activity during decision-making. This effect was not observed in the aIns for any behavioral measures (i.e., HE, HPE, HME choices, or kEp) (Fig. S5-1), highlighting a region-specific modulation by dmPFC/dACC lactate.

**Figure 5:**
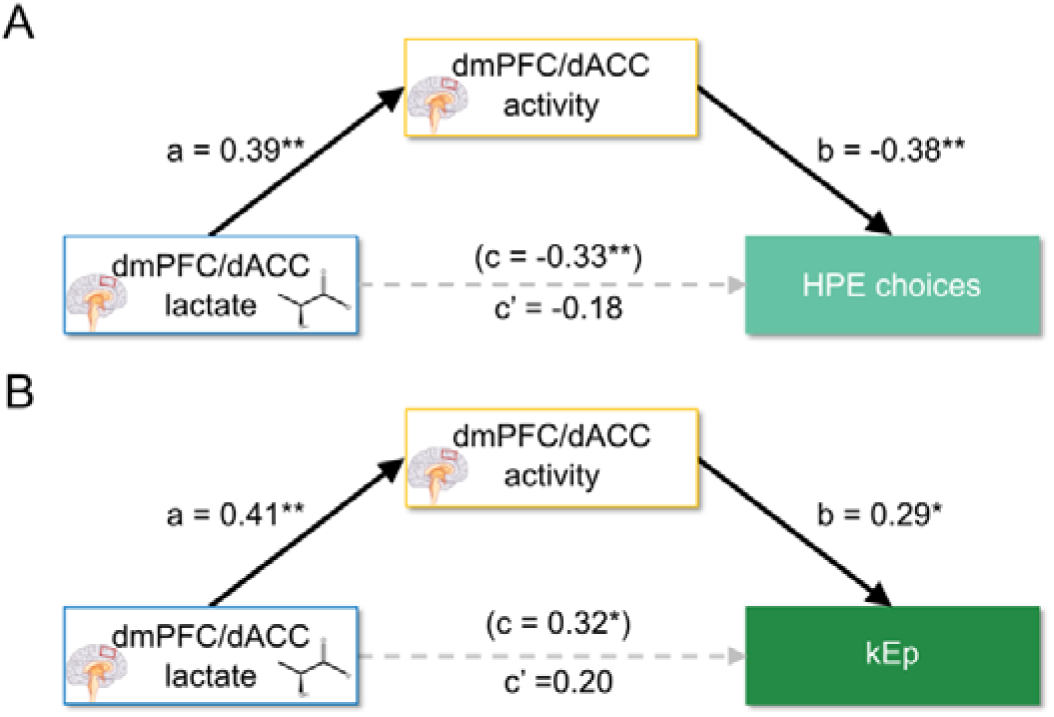
Mediation analysis of dmPFC/dACC lactate on motivated behavior via dmPFC/dACC activity. Significant paths are depicted with continuous black lines, while non-significant paths are represented with dotted grey lines. Path c represents the coefficient for the direct path before considering the mediator, while path c’ represents the coefficient for the direct path after considering the mediator in the same equation, and path a and b are the coefficients for the indirect pathway (i.e. the mediation). The equivalent tests for the aIns can be found in **Fig. S5-1** . Significance levels for each path: ***p<0.001; **p<0.01; *p<0.05. **A**] High physical effort (HPE) choices: a mediation analysis shows dmPFC/dACC lactate levels influencing HPE choices mediated by dmPFC/dACC fMRI regression estimate for Ech (N = 60). **C**] Sensitivity to physical effort kEp: Path analysis shows dmPFC/dACC lactate levels influencing kEp, mediated by dmPFC/dACC fMRI regression estimate for Ech (N = 57).

Expanding our analysis to incorporate plasma lactate, structural equation modeling (SEM) was used to assess its impact alongside brain lactate levels on decision-making and neural correlates. This broader model confirmed the linkage between plasma lactate and dmPFC/dACC lactate levels and supported the role of dmPFC/dACC lactate in driving both motivated behavior and neural activity during HPE decisions (Fig. 6A). Again, similar results were obtained for kEp, reinforcing the idea that lactate’s influence on HPE choices might be mostly driven by its specific impact on sensitivity to physical effort (Fig. 6B). Although the explained variance for our models may seem modest for each of the components considered (0.1 ≤ R² ≤ 0.3), these values are consistent with the expected range for neuroimaging studies, which typically explain less than 20% of inter-individual behavioral variance [68, 69]. It is noteworthy to consider that a single neurometabolic measure (lactate) taken in one single brain area (the dmPFC/dACC) two hours before the actual task explains such a proportion of neural and behavioral variance.

**Figure 6:**
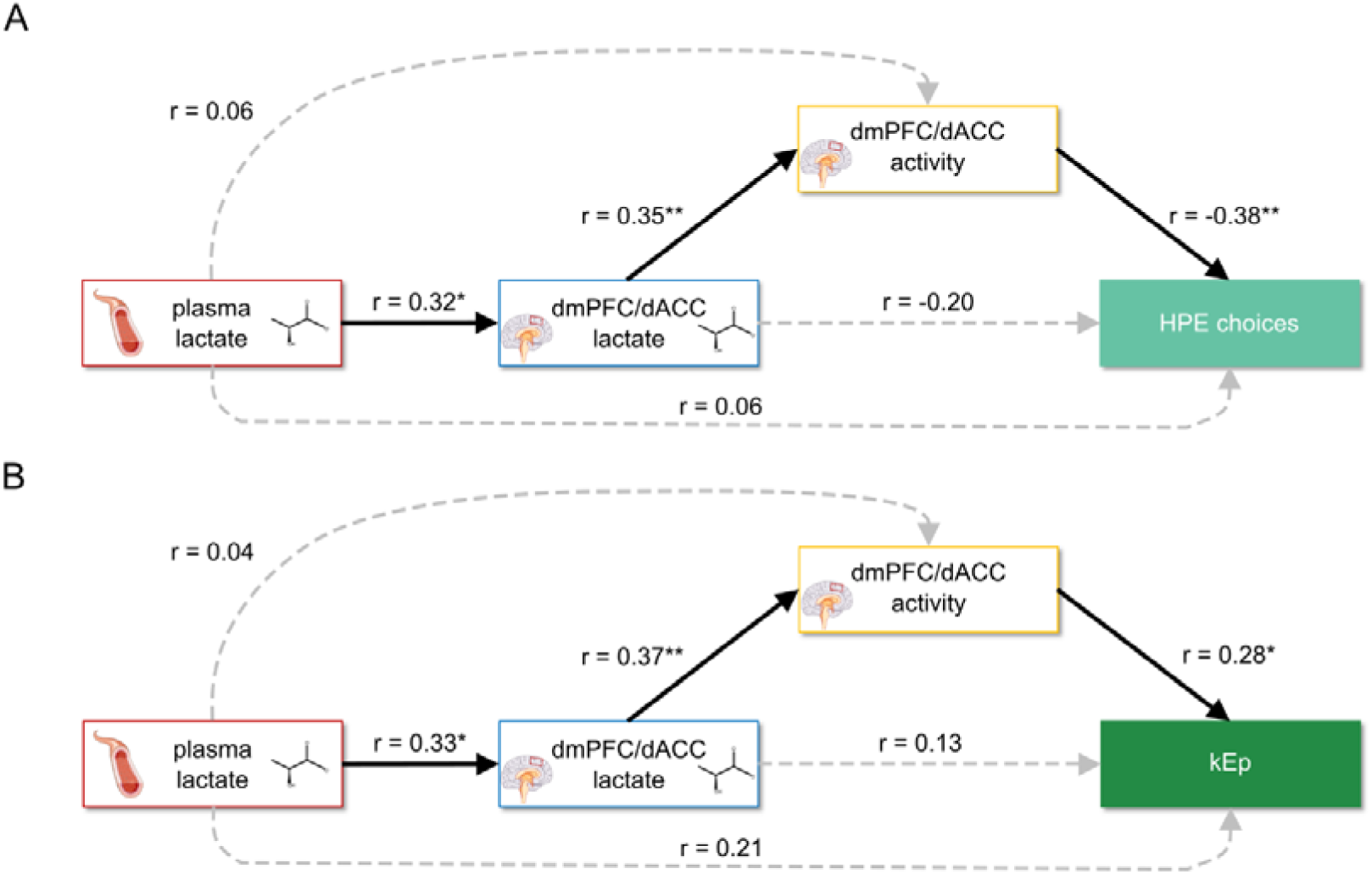
Structural Equation Modeling of dmPFC/dACC lactate influence on motivated behavior via dmPFC/dACC activity, including plasma lactate. Significant paths are depicted with continuous black lines, while non-significant paths are represented with dotted grey lines. Significance levels for each path: ***p<0.001; **p<0.01; *p<0.05. A] Path analysis for high physical effort (HPE) choices: Examining the influence of plasma lactate on HPE choices, through dmPFC/dACC lactate and activity (N = 59). The model explained variance for each of the parameters of the path analysis was the following: dmPFC/dACC lactate R² = 0.099, dmPFC/dACC Ech estimate R² = 0.138, HPE choices R² = 0.226. B] Path analysis for the sensitivity to physical effort kEp: Examining the influence of plasma lactate on kEp, through dmPFC/dACC lactate and activity (N = 56). The model explained variance for each of the parameters of the path analysis was the following: dmPFC/dACC lactate R² = 0.106, dmPFC/dACC Ech estimate R² = 0.151, kEp parameter R² = 0.208.

In summary, our path analyses suggest that plasma lactate levels are associated with dmPFC/dACC lactate, and it is the latter that modulates decisions and sensitivity regarding physical effort, primarily through its influence on dmPFC/dACC neural activity. These findings provide insights into the mechanistic pathways through which metabolic states may influence motivational processes, particularly in contexts requiring significant physical effort.

## Discussion

Despite the well-documented roles of the dmPFC/dACC and aIns in regulating goal-directed behavior [8, 51, 64], the mechanisms underlying individual variability in the willingness to exert effort remain largely unexplored. Our findings indicate that dmPFC/dACC lactate levels explain individual differences in decision-making processes related to physical effort, primarily through their impact on neural activity within this region. By integrating metabolic profiling, functional neuroimaging, behavioral assays, and computational modeling, our study suggests that while plasma lactate levels may contribute to dmPFC/dACC lactate levels, it is the latter that modulates decisions requiring physical effort through its influence on neural activity. Thus, our study supports the role of lactate as a critical modulator within the dmPFC/dACC, particularly under conditions that demand physical exertion. Challenging the view of lactate as a booster of physical effort [70] and an antidepressant [71–73], our results instead emphasize its involvement in modulating neural activity [74], ultimately dampening physical effort-based decision-making behaviors at high lactate levels. This novel metabolic-neural pathway deepens our understanding of the neurobiological underpinnings of motivation and may open new avenues for clinical intervention.

Notably, our study observed that neural activity in the dmPFC/dACC escalated with the level of effort chosen, supporting its role as an energization signal [50, 55] rather than merely indicating the costs of effort to other brain areas [8, 62]. This increase in neural activity correlated with elevated lactate levels, which is consistent with lactate’s capacity to modulate neuronal excitability through various receptors and channels [74–76]. Additionally, the steepness of the dmPFC/dACC BOLD response, as a function of chosen effort difficulty, inversely correlated with subjects’ propensity to select high-effort options, as previously observed [50]. This suggests that increased dmPFC/dACC and aIns activity reflect an increased perception of effort costs [67], hinting at a cost signal that modulates willingness for exertion. Supported by lesion and stimulation studies indicating a causal role of the dmPFC/dACC in effort-based decision-making [55, 77, 78], our findings suggest a mechanism where increased neural recruitment, influenced by lactate, serves not only as an energizing signal but also carries a neural cost critical in shaping decisions related to effort expenditure.

Our findings introduce the compelling possibility that plasma lactate levels, particularly elevated under energetically demanding conditions such as exercise, affect dmPFC/dACC lactate levels, thereby impacting brain function and cognitive processes [22–24]. The ability of lactate to cross the blood-brain-barrier via specialized monocarboxylate transporters [79, 80] allows it to serve not just as a metabolic fuel [81], but also as a signaling molecule [35, 36] that informs the brain about the body’s overall metabolic state. According to our findings, this signaling role of lactate might be particularly relevant in situations of high energy demands. For example, lactate’s signaling could be particularly important under exercise [82] or pathological states such as inflammation, mitochondrial dysfunction or brain lesions [83–85]. Although we did not measure dynamic changes in lactate levels to directly link with task-induced fatigue, and no association was found between baseline levels and subjective fatigue as measured via questionnaires, the signaling role of peripheral lactate could partially explain the emergence of central fatigue [86], a common feature across these conditions, and it could therefore serve as a potential mechanism for preserving energetic resources. Additionally, the well-documented correlation between elevated blood lactate levels and subjective ratings of physical effort during exercise [87–89] supports this signaling role. Recent studies also show that injections leading to elevated lactate levels lead to decreased locomotor activity [86, 90, 91] which, in line with our findings, suggest that elevated plasma lactate levels may serve as a signal to the dmPFC/dACC, potentially discouraging the selection of tasks that require significant physical effort. However, peripheral administration’s effects on subjective fatigue and effort perception [92, 93] or in effort performance and locomotor activity [72, 73] are not always consistent, indicating a need for further well-powered interventive studies to clarify lactate’s role in effort-based decision-making and fatigue. An interaction with other plasma metabolites may be crucial for lactate’s signaling effect [94, 95], as high lactate levels combined with ATP and H^+^ in muscle can induce sensations of peripheral fatigue and pain, whereas none of these metabolites alone produce such effects [93]. Of note, the fact that our data suggest that lactate acts as a signaling molecule regulating the willingness to exert physical efforts does not preclude 1) that, as ketone bodies, lactate can also be used as an alternative fuel to glucose by the brain, especially during mental and physical exercise [81, 96–98] and 2) that exercise-derived lactate, in addition with other so-called ‘exerkines’, could bear a mid/long-term positive influence on physical and mental health [99–101].

While the specificity of our results regarding the dmPFC/dACC merits attention, caution is warranted due in part to the smaller number of participants with valid aIns ^1^H-MRS lactate measurements, which was additionally always performed after the dmPFC/dACC. and the fact that lactate levels in the dmPFC/dACC and aIns showed a moderate trend to correlate. Plasma lactate concentrations correlated specifically with dmPFC/dACC, but not with aIns, lactate levels. This variability aligns with documented significant regional differences in lactate concentrations across the resting-state human brain [49], as well as marked metabolic differences between brain regions [102–104]. Specifically, the dmPFC/dACC is particularly notable for its high levels of energy consumption and aerobic glycolysis [102, 104, 105]—a process that not only supports high energy demands but also produces lactate as a by-product. Variations in blood-brain barrier permeability, metabolic activity, vascularization, and the expression and regulation of lactate transporters across brain different regions also contribute to the differential distribution and utilization of lactate.

Addressing our study’s limitations, we acknowledge that the explained variance by dmPFC/dACC activity and lactate levels is partial, suggesting the involvement of additional neural circuits and metabolic pathways in effort-based decision-making. Future studies should explore other regions implicated in motivation, such as the ventral striatum and dorsolateral prefrontal cortex [106–109], which also correlated with the difficulty of chosen effort across our two tasks. Investigating other metabolites alongside lactate, could elucidate the complex interactions underpinning effort-based decision-making [12–14]. Notably, our findings reveal no significant correlations between lactate levels in plasma, dmPFC/dACC, or aIns and motivation for mental effort, despite the established role of the dmPFC/dACC in governing mental effort engagement in such tasks [55, 65, 110, 111]. However, our results cannot rule out that dmPFC/dACC lactate is also involved in other components of motivation. Furthermore, a multivariate analysis of the same dataset indicates that a combination of aspartate, glutamate, and lactate in the dmPFC/dACC can predict inter-individual differences in mental-effort-based decision-making [112], suggesting that lactate is also relevant for mental effort computations but its impact on mental effort may depend on its interaction with other metabolites. These findings underscore the complexity of metabolic contributions to cognitive processes and highlight the need for further investigation. Future studies, potentially involving animal models, are warranted to dissect the causal mechanisms by which lactate influences effort-related neural activity.

In conclusion, our study supports the crucial role of lactate in the dmPFC/dACC as a metabolic signal that regulates motivation for physical effort, offering new insights into the interplay between metabolic states and cognitive functions. This work enriches our understanding of motivation, supporting the view that metabolic states intricately influence cognitive functions and decision-making. These findings emphasize the need for an integrated approach that considers both neurobiological and metabolic mechanisms to fully understand the complexities of human motivation and decision-making.

## Materials & Methods

### Experimental Design

The current study is part of a larger experimental cohort [112]. In brief, first, participants were brought to a medical center where professional nurses collected their blood (Fig. 1A) which served to measure lactate levels in the plasma. We then acquired participants’ proton magnetic resonance spectroscopy (^1^H-MRS) metabolites in the dmPFC/dACC and in the aIns while participants were resting in the MRI scanner (Fig. 1A, Fig. S1-2A) providing dmPFC/dACC and aIns lactate levels. Then, participants performed extensive training and calibration out of the scanner. This training was essential to familiarize participants with the tasks and to calibrate monetary incentives and effort levels. Finally, participants went back in the scanner where they performed the behavioral task during four fMRI blocks alternating between physical and mental effort blocks (Fig. 1A). The behavioral task was an effort-based decision-making task allowing to assess how people weigh monetary incentives (rewards and punishments) and efforts (physical and mental). On each trial, participants had to perform the selected effort to obtain the associated reward (or avoid increasing the associated loss). The physical effort consisted in squeezing a handgrip dynamometer for variable durations (from 0.5 to 4.5s) but always with the same strength threshold (55% of their maximal voluntary contraction force) (Fig. 1B). The mental effort consisted in providing a certain number of correct answers (based on individual calibration) within a 2-back task with a fixed duration of 10s in all trials (Fig. 1C). Participants’ final reward depended partly on their performance in the behavioral task.

### Participants

In total, 75 right-handed healthy volunteers (N = 40 females) participated in this study which was approved by the Cantonal Ethics Committee of Vaud (CER-VD), Switzerland. The number of volunteers was defined to capture a moderate expected correlation (r ∈ [0.4-0.6]) between dmPFC/dACC lactate and physical effort sensitivity based on a calculation in G*Power 3.1.9.7 (N.subjects ∈ [25–63]), in agreement with a previous study which showed a moderate relationship (|r| = 0.5) between ventral striatum metabolites and effort perception [12]. Participants were recruited through the Université de Lausanne (UNIL) LABEX platform, and through online and printed announcements in the city of Lausanne. To be involved in the study, participants had to speak French fluently and be aged between 25 and 40 years old. They were also screened for exclusion criteria: regular use of drugs or medications, history of neurological disorders, and contraindications to MRI scanning (pregnancy, claustrophobia, recent tattoo near the neck, metallic implants). All participants gave their signed informed consent prior to participation in the study. Before coming to the lab, participants filled out the inclusion/exclusion criteria and several online behavioral questionnaires using the online Qualtrics platform (Qualtrics, Provo, Utah, USA). To ensure a wide range of motivational profiles in the healthy population sampled, we used the self-rated Montgomery Asberg Depression Rating Scale (MADRS-S) questionnaire for depression [113] with a lenient cut-off of 4 [114], as studying depression was not the aim of the study. We therefore ensured to have both individuals with a low score in the MADRS-S questionnaire (MADRS-S < 4, N = 37) and individuals with mild/high scores (MADRS-S ≥ 4, N = 38). Four of the participants did not perform the behavioral experiment in the MRI due to technical issues (two did the behavior out of the MRI and two decided to quit the experiment before). Two more subjects were excluded because of a lack of whole-brain coverage in the fMRI acquisition. Six more subjects selected the high effort (HE) option in more than 95% of the trials in the two blocks of one of the two tasks, making it impossible to fit their behavior. After removing these twelve subjects, we were left with N = 63 subjects (see Table S1 for demographic details). For some of the analysis, we also had to remove a few more subjects due to plasma (N = 1), dmPFC/dACC (N = 2) or aIns (N = 18) lactate values missing. Participants were paid 70 CHF for completing the task and 10 CHF per hour spent in the experiment. In addition, they were given a fixed amount of 4 CHF for each time they performed a physical or mental maximal performance although they were told that the payoff depended on their performance on these trials to ensure that they calibrated the task properly. Finally, they were also given a bonus related to their choices and performance during the indifference point measurement and in the main task. On average, participants obtained 203.17 ± 2.28 CHF for performing the task.

Before coming to the laboratory, participants filled out behavioral questionnaires, including the MADRS-S, online. The day before the experiment, they were instructed to refrain from performing any intense physical activity (sports). The experiment was always starting around 2pm, participants were asked to refrain themselves from eating and drinking anything else than water (particularly no sweet or acid drinks, or drinks containing caffeine) and to not smoke for at least an hour before the start of the experiment. These instructions were given in order to minimize as much as possible any bias on the metabolic measures of the blood and of the brain.

### Behavioral Task

#### General overview

The task was designed and run using Psychtoolbox 3 [115], and implemented in Matlab (The Mathworks Inc., USA). We first calibrated the levels of effort to the individual’s capacity and monetary incentives based on a short indifference point calibration (see Supplementary Methods). Then, subjects got to practice the different levels of effort and a few choices to know what they would be exposed to in the main task in the scanner. The main task in the fMRI scanner consisted of performing 4 blocks of effort-based decision-making and performance tasks. Blocks were decomposed in 2 mental and 2 physical effort blocks that were always alternating with each other (Fig. 1A). The order of the physical and mental blocks was counterbalanced across participants during both training and fMRI sessions. During a given block, participants first performed two maximal physical (or mental) effort capacity trials, then they accomplished 54 choice trials always immediately followed by the associated effort (Fig. 1B-C), and at the end of a block, they again had to perform two maximal physical (or mental) capacity trials.

#### The behavioral task

The task consisted of 4 fMRI blocks. These blocks were organized into two physical and two mental effort blocks which were alternating (Fig. 1A). The order was counterbalanced across participants. Each block started and finished with two measures of participants’ maximal capacity. Each block was composed of 54 trials, each divided into a choice and an effort period. The choice period consisted in selecting between two options, each associated to different levels of monetary incentives and effort. The effort period consisted in exerting a physical or a mental effort corresponding to the effort level they had selected with performance progression indicated through a yellow pie chart disappearing as performance increased. In total, participants completed 216 trials. We informed participants that they should be able to do all types of effort in all trials so that their choices should depend on their subjective preferences rather than on their estimation of being capable to do the task (i.e. risk discounting). To further minimize the influence of risk discounting on subjective preferences, participants were informed that even if they couldn’t reach the end of the pie chart for a given trial, they would still obtain a proportion of the reward at stake (or avoid a proportion of the punishment at stake respectively). Before and after each block, participants had to perform twice their maximal effort capacity as during the calibration period. In the physical effort task, this consisted in squeezing the grip as hard as possible within 5 seconds. In the mental effort task, this consisted in giving as many correct answers as possible in the 2-back task, within 10 seconds. This step aimed at assessing how effort capacity varied over time, especially regarding the possible reduction in effort capacity due to fatigue. To ensure that participants were complying we told them that they would be rewarded based on their performance every time their maximal performance would be required, although they always got a fixed amount of 4 CHF for each series of maximal performance attempts.

#### Choice period

During the choice period, participants were presented with two options (Fig. 1B-C), one fixed low incentive/low effort (+0.5 CHF for rewards or -0.5 CHF for punishments and effort level 0), and one high incentive/high effort, which varied independently in effort and incentive level. Each effort level was represented by a yellow pie chart, with bigger yellow slices indicating more difficult efforts. Monetary incentives were composed of 8 levels (4 rewards and 4 punishments) and effort difficulty was composed of 4 levels for both tasks. The left/right position on the screen of the two options was counterbalanced across trials. On each trial, participants used a four-buttons response pad (Current Designs Inc., Groningen, Netherlands) to select an option (left choice with the 2 left buttons/right choice with the 2 right buttons) and also to indicate how confident they were that this was the best choice (low confidence had to be indicated with the 2 buttons in the middle and high confidence with the 2 buttons at the extremes of the response pad). In case participants did not make a choice, the low incentive/low effort was chosen by default, and we discarded this trial from the analysis. After their choice selection, the selected option was displayed, framed by continuous (for high confidence) or dotted (for low confidence) lines reflecting the level of confidence.

#### Effort period

At the onset of the effort period, a yellow pie chart, representing the amount of effort still required to fulfill the trial, appeared at the center of the screen. To obtain the reward (or avoid doubling the loss) that they selected, the objective of the participants was to make that yellow pie chart disappear by applying sufficient effort in the allocated time.

During the physical effort phase, participants had to squeeze the handgrip for the selected duration, at least at 55% of their MVC within 6s (Fig. 1B). A vertical bar representing participants’ real-time force was displayed to the left, superimposed by a red bar representing the required threshold to exceed (55% of their MVC). If participants exerted strength above that threshold, the yellow pie chart diminished linearly over time until it disappeared, ending the effort phase, otherwise, the yellow pie chart was staying frozen, until participants reached the threshold.

During the mental effort period, participants had to provide a determined number of corrected answers within 10 seconds (Fig. 1C). Each correct response removed a portion of the yellow pie chart, until performance was achieved, or the time run out. To prevent participants answering randomly to win speed over accuracy, the yellow pie chart would increase in size every time participants gave an incorrect answer, unless the pie chart was still in its initial stage of completion when the wrong answer was provided, in which case it remained unchanged.

Participants were informed that even if they could not complete the requested amount of effort in the correct amount of time for a given trial (i.e. the yellow circle was still visible at the end of the trial), they would still obtain a proportion of the reward at stake, corresponding to the percentage of effort performed. At the end of the effort phase, feedback on the amount of money lost or won was displayed on the screen for each trial.

### Behavioral Analysis

All data were analyzed using MATLAB 2021a (The MathWorks). Participants’ choices were fitted with a softmax model using Matlab’s VBA toolbox (https://mbb-team.github.io/VBA-toolbox/) which implements Variational Bayesian analysis under the Laplace approximation [116]. This model allowed us to extract behavioral parameters reflecting the sensitivity to monetary incentives (kR, kP), to physical and mental effort (kEp, kEm), along with other temporal effects (kFp, kLm) and a general bias towards selecting the high or the low effort option (kBias). Details of the modeling approach can be found in the Supplementary Methods.

### Statistics

The results reported in the main text, unless specified otherwise, correspond to the mean ± standard error of the mean. For all the correlations reported between variables, we also removed any subject who had a value above or below three standard deviations away from the median on any of the measurements. When testing the specificity of our results, correlation coefficients were compared using the R cocor package [117] with a one-tailed Steiger’s test [118] at α = 0.05 and confidence level = 0.95 between two dependent (because variables were coming from the same individuals) overlapping (when a variable was used in both correlations) groups.

### Plasma Acquisition

Blood was extracted at the moment participants arrived for the experiment, just after they signed the consent forms (Fig. 1A). Professional nurses from the Point Santé of the École Polytechnique Fédérale de Lausanne (EPFL) or from the Arcades medical center extracted the blood from participants in S-monovette 1.2 mL tubes (Sarstedt, Nümbrecht, Germany) containing EDTA K2E to avoid blood clotting. Blood was immediately stored in ice at 4°C and transported to our laboratory within approximately 15 minutes. Then, blood tubes were centrifuged at 1100g and 4°C for 15 minutes in a PK 120 R ALC centrifuge to dissociate the plasma from blood cells. Finally, plasma was extracted in tubes of 100μL in which 5μL of protease inhibitor cocktail was added. Samples were subsequently stored at -80°C until further analysis.

### Plasma Lactate Analysis

Plasma lactate concentrations were measured using a liquid chromatography instrument (Waters Aquity, Milford, MA, USA) coupled to a tandem mass spectrometer (Sciex 6500+, Toronto, Canada) allowing to perform liquid chromatography coupled to mass spectrometry (LC-MS). Following Dei Cas and colleagues [119], 40μL of plasma samples were deproteinized and then derivatized with 3-nitrophenylhydrazine (3-NPH). Derivatized compounds were then quantified by reversed-phase liquid chromatography on a Raptor ARC-18 UHPLC column (2.1 x 100 mm; 1.7µm) and analytes were detected in negative mode using multiple reaction monitoring (MRM) transitions specific for each analyte. The concentration of lactate was then calculated by comparison of the ratios between the signal intensity of lactate to labeled lactate used as internal standard, via its corresponding calibration curve. Except one subject where there was not enough plasma to perform the analysis, all plasma lactate levels were within the range of detection and were therefore included in the analyses.

### 1*H-MRS* Data Acquisition

Proton magnetic resonance spectroscopy (^1^H-MRS) was acquired in a 7 Tesla scanner (Magnetom, Siemens Medical Solutions, Erlangen, Germany) equipped with a single channel quadrature transmitter and a 32-channel receive coil (Nova Medical Inc., MA, USA). Shimming was performed using FAST(EST)MAP [120] for both first- and second-order to optimize magnetic field homogeneity. Head movements were minimized by positioning foam pieces around the participants’ heads. In addition, OVS bands were carefully placed to suppress signals from outside the voxel-of-interest (VOI), in particular extra-cerebral fat signals. The voxels were positioned using T1-weighted magnetization prepared 2 rapid gradient echo (MP2RAGE) [121] with the following acquisition parameters (TE = 1.88ms; TR = 6s, TI1/TI2 = 800/2700 ms, α1/α2 = 7°/5°, slice thickness = 1 mm, FOV = 192 × 192 × 192 mm^3^, matrix size = 192 × 192 × 192, bandwidth = 240 Hz/Px). We always acquired the dmPFC/dACC voxel first, followed by the left aIns voxel.

The dmPFC/dACC voxel (20 x 20 x 20 mm^3^) was positioned using MP2RAGE anatomical images, by aligning it on the midline of the axial and coronal planes. In the sagittal plane, the voxel horizontal borders were defined to be parallel to the cingulate sulcus. The anterior vertical border was defined to be above the genu of the corpus callosum.

The left aIns voxel (20 x 20 x 20 mm^3^) was aligned to the superior and to the anterior peri-insular sulcus [122] on the sagittal plane. On the axial and coronal plane, we moved the voxel to maximize the signal quality and the number of voxels overlapping the insula, while minimizing contact with the ventral pallidum.

Semi-adiabatic spin echo full intensity acquired localized (sSPECIAL) [123–125] was used to acquire 50 MR spectra blocks (2 average/block) per region of interest, with the following acquisition parameters (TE = 16 ms; TR = 5 s, spectral bandwidth 4000 Hz, and a number of points of 2048). Outer Volume Suppression (OVS) bands were positioned around the VOI to minimize spectral contamination from lipid and water signals originating from peripheral regions around the brain.

### 1*H-MRS* Data Analysis

All spectra were corrected for frequency/phase shifts and averaged. Then, we used LCModel for spectral fitting and lactate quantification [126], with a basis set that included simulated metabolite spectra and an experimentally measured macromolecule baseline (Fig. S1-2B). In addition, a spectrum of unsuppressed water was acquired and used as an internal reference for absolute lactate quantification in LCModel. Furthermore, following lactate quantification in LCModel and correction, any computed metabolite concentrations with a Cramér-Rao lower bounds (CRLB) higher than 50 % were excluded. This led us to remove 2 subjects for the analyses including dmPFC/dACC lactate and 18 subjects for aIns lactate.

Anatomical T1-weighted images were used to compute the tissue composition within each MRS voxel. The T1-weighted images were segmented into grey matter (GM), white matter (WM), and cerebrospinal fluid (CSF) in SPM12 toolbox (Wellcome Trust Center for NeuroImaging, London, UK) with the Marsbar toolbox (https://marsbar-toolbox.github.io/) working in Matlab (The Mathworks, US). Metabolite concentrations in each MRS voxel were corrected for the CSF fraction, assuming water concentrations of 43,300 mM in GM, 35,880 mM in the WM, and 55,556 mM in the CSF. The dmPFC/dACC and aIns spectrum resulted in an overall signal-to-noise ratio (SNR) LCModel output and FWHM of 88 ± 12, 0.029 ± 0.028 ppm, and 53 ± 17, 0.036 ± 0.01 ppm, respectively. The average CRLB for lactate was of 0.076 ± 0.003 in the 61 remaining dmPFC/dACC subjects and of 0.146 ± 0.017 in the 45 remaining aIns subjects, confirming the good quality of the data in the subjects included in the analysis. The quality of the ^1^H-MRS data was also confirmed by the observation of the characteristic lactate doublet at ∼1.33ppm (Fig. S1-2B) when looking at the MRS spectra.

### fMRI Data Acquisition

Functional and structural brain imaging data was collected using a Siemens Magnetom 7T scanner (Siemens, Erlangen, Germany) equipped with a 32 channel Head/Neck coil (Nova Medical Inc., MA, USA). Structural T1-weighted images were co-registered to the mean echo planar image (EPI), segmented and normalized to the standard T1 MNI template and then averaged across subjects for anatomical localization of group-level functional activations. Functional T2*-weighted EPIs were acquired with BOLD contrast using the following parameters: repetition time TR: 2.00 seconds, echo time TE = 26 ms; flip angle = 63°; number of slices = 63; slice thickness = 2.00 mm; field of view = 304 mm; multiband acceleration factor = 3. Note that the number of volumes per block was not predefined because all responses were self-paced. Volume acquisition was just stopped when the task was completed.

### fMRI Data Analysis

Functional MRI data were preprocessed and analyzed with the SPM12 toolbox (Wellcome Trust Center for NeuroImaging, London, UK) running in Matlab 2021a. Preprocessing consisted of spatial realignment, normalization using the same transformation as anatomical images, and spatial smoothing using a Gaussian kernel with a full width at a half-maximum of 8 mm.

Preprocessed data were analyzed with a standard general linear model (GLM) approach at the first (individual) level and then tested for significance at the second (group) level. All GLM included six movement regressors generated during the realignment procedure. First-level images were masked with an inclusive mask derived from the group of subjects included in the analysis. The mask was built by first creating a first level grey matter mask for each subject through SPM’s segmentation procedure during preprocessing. These individual masks were then averaged across subjects and only voxels with a probability higher than 5% to be in the grey matter were kept. Individual fMRI blocks where subjects picked up one option (either the high effort or the low effort) more than 94% of the time were removed from the analysis for all GLMs. Moreover, for GLM4 (see Supplementary Methods), we also removed any fMRI block where high effort or low effort choices were always associated to a single high effort level, since SPM could not compute a regression otherwise.

Our first GLM (GLM1) included one categorical regressor to model fixation crosses onsets, a second categorical regressor to model the choice onset, which was parametrically modulated by the effort chosen (Ech), the subjective value of the chosen option (SVch) derived from our computational model and choice deliberation time (DT), another regressor modeled the onset of the display of the chosen option, another regressor modeled the onset of the effort period and a last regressor modeled the feedback onset. All categorical regressors were modeled with a stick function. All parametric modulators were zscored per block. The parametric modulators were not orthogonalized, so that they were competing to explain the variance of the BOLD signal.

For region-of-interest (ROI) analyses, we employed ^1^H-MRS-derived masks for the dmPFC/dACC and the aIns ensuring the analysis of fMRI data without double dipping [127] and that the fMRI activations analyzed correspond to the exact same location where ^1^H-MRS was measured. To create these masks, we first extracted the coordinates of the individual ^1^H-MRS voxels in native-space. Then we applied the same transformation to these voxels as the one applied to fMRI images based on co-registration and we obtained the corresponding voxel coordinates in MNI space for each individual. The density map of the voxels covered across all individuals in MNI space can be found in Figure 3B and in Figure S1-2A. The activity of each region-of-interest (ROI) was extracted as the average level of activity of all the fMRI voxels located inside the individual ^1^H-MRS mask. Subjects where ^1^H-MRS could not be measured for a given ROI were subsequently also excluded from the ROI analysis of fMRI. Violin plots with ROI fMRI results were generated using the violinplot Matlab function developed by Bastian Bechtold (https://github.com/bastibe/Violinplot-Matlab, doi: 10.5281/zenodo.4559847).

### Overlap between ^1^H-MRS voxel and fMRI clusters

The overlap between ^1^H-MRS voxels and fMRI Ech clusters was performed by extracting the dmPFC/dACC fMRI cluster for Ech (at a voxel-wise threshold of p < 0.001 with a family-wise error (FWE) correction for multiple comparisons, N = 63) and the left aIns fMRI cluster for Ech (at a voxel-wise threshold of p < 0.05 with a FWE correction for multiple comparisons, N = 63) and the ^1^H-MRS dmPFC/dACC (N = 63) and aIns (N = 60) voxels sampled across all the 63 participants included in MNI space and then to look at the percentage of overlap between the two. Any voxel sampled with ^1^H-MRS in at least one subject and as well present in the average fMRI cluster for Ech would then be counted as an overlap. We also performed a second test by restricting the analysis to the ^1^H-MRS voxels sampled in at least 90% of the participants and extracted the overlap accordingly.

### Structural Equation Modeling (SEM) Analysis

The SEM analysis was performed using R Statistical Software (version 4.3.2; R Core Team 2023; https://www.r-project.org/) running in RStudio (version 2023.12.1+402; RStudio Team (2020); http://www.rstudio.com/) with the lavaan (v0.6.16) package [128]. Before launching the mediation and SEM analysis, we filtered any subject who had a missing value on any of the measurements (between plasma levels of lactate, brain levels of lactate, fMRI regression estimate, or behavioral output) and we also removed any subject who had a value above or below three standard deviations away from the median on any of the measurements involved in the analysis. Note that this was done independently for each analysis therefore they do not always include the same number of subjects (N = 41-43 for aIns analyses and N = 56-59 for dmPFC/dACC analyses).

### Data And Code Availability

The original code developed for the task has been deposited at https://github.com/NicolasClairis/physical_and_mental_E_task.git and is publicly available as of the date of publication. The second-level whole-brain fMRI maps have been uploaded in Neurovault and can be found at: https://neurovault.org/collections/17029/. Any additional information required to reanalyze the data reported in this paper is available from the lead contact upon reasonable request.

## Acknowledgements

We would like to thank Mathias Pessiglione, Antonius Wiehler, Alizée Lopez-Persem, Bogdan Draganski for their contributions to the design of the experiment. We would also like to thank the Center for Biomedical Imaging (CIBM) staff for their help in fMRI data acquisition, in particular Sandra Da Costa. We thank the nurses from the École Polytechnique Fédérale de Lausanne (EPFL) Point Santé (Viviane Depuydt-Linder and Chiyama Mathivathanasekaram), the personnel from the Centre Médical des Arcades and the members of the Laboratory of Behavioral Genetics (LGC) for their support in blood collection and processing. We would also like to thank Antoine Lutti for assistance in the fMRI analysis and Bernard Cuenoud (Nestlé Health Sciences, Lausanne), Jean-Philippe Godin [Nestlé Research (NR), Lausanne], Stefan Christen (NR), Karine Meisser (NR), Olivier Ciclet (NR), Adrien Frezal (NR), and Irina Monnard (NR) for their valuable contributions to the plasma lactate analyses. We also thank Simone Astori for his help on the design of the figures and Martin Picard for his valuable advice, which greatly contributed to the improvement of the manuscript.

## Funding

NC has received funding from the European Union’s Horizon 2020 research and innovation program under the Marie Sklodowska-Curie grant agreement N°101032219, the Foundation for the encouragement of Nutrition Research in Switzerland (SFEFS) under the grant agreement n° 607 and the Novartis Foundation for medical-biological Research under the grant agreement n°#21B110. The project was also funded by intra-mural funding from the École Polytechnique Fédérale de Lausanne (EPFL) to C.S..

## Author Contributions

Conceptualization: AB, NC, CS

Methodology: AB, JB, NC, CS

Software: AB, NC, LX

Formal Analysis: AB, NC

Investigation: AB, NC

Resources: CS, LX

Writing-original draft: AB, NC, CS

Visualization: NC

Supervision: CS, LX

Project Administration: CS

Funding Acquisition: NC, CS

## Conflict of Interest

C.S. is a member of the scientific advisory board of Amazentis/Timeline S.A and Vandria S.A.

## Notes

https://neurovault.org/collections/17029/

https://github.com/NicolasClairis/physical_and_mental_E_task

### Summary of Updates

We fixed some small omissions and typos and incorporated reviewers' feedback especially regarding claims of specificity which were not properly tested in the previous version of the manuscript.

## References

1. An H-Y, Chen W, Wang C-W, Yang H-F, Huang W-T, Fan S-Y. The Relationships between Physical Activity and Life Satisfaction and Happiness among Young, Middle-Aged, and Older Adults. Int J Environ Res Public Health. 2020;17:4817.

2. Bernacer J, Martinez-Valbuena I, Martinez M, Pujol N, Luis E, Ramirez-Castillo D, et al. Brain correlates of the intrinsic subjective cost of effort in sedentary volunteers. Prog Brain Res. 2016;229:103–123.

3. Cillekens B, Lang M, van Mechelen W, Verhagen E, Huysmans MA, Holtermann A, et al. How does occupational physical activity influence health? An umbrella review of 23 health outcomes across 158 observational studies. Br J Sports Med. 2020;54:1474–1481.

4. Bonnelle V, Veromann K-R, Burnett Heyes S, Lo Sterzo E, Manohar S, Husain M. Characterization of reward and effort mechanisms in apathy. J Physiol Paris. 2015;109:16–26.

5. Chong TT-J, Bonnelle V, Husain M. Chapter 4 - Quantifying motivation with effort-based decision-making paradigms in health and disease. In: Studer B, Knecht S, editors. Progress in Brain Research, vol. 229, Elsevier; 2016. p. 71–100.

6. Le Heron C, Apps M a. J, Husain M. The anatomy of apathy: A neurocognitive framework for amotivated behaviour. Neuropsychologia. 2018;118:54–67.

7. Lopez-Gamundi P, Yao Y-W, Chong TT-J, Heekeren HR, Mas-Herrero E, Marco-Pallarés J. The neural basis of effort valuation: A meta-analysis of functional magnetic resonance imaging studies. Neurosci Biobehav Rev. 2021;131:1275–1287.

8. Pessiglione M, Vinckier F, Bouret S, Daunizeau J, Le Bouc R. Why not try harder? Computational approach to motivation deficits in neuro-psychiatric diseases. Brain. 2018;141:629–650.

9. Morella IM, Brambilla R, Morè L. Emerging roles of brain metabolism in cognitive impairment and neuropsychiatric disorders. Neuroscience & Biobehavioral Reviews. 2022;142:104892.

10. Ülgen DH, Ruigrok SR, Sandi C. Powering the social brain: Mitochondria in social behaviour. Current Opinion in Neurobiology. 2023;79:102675.

11. Yellen G. Fueling thought: Management of glycolysis and oxidative phosphorylation in neuronal metabolism. Journal of Cell Biology. 2018;217:2235–2246.

12. Strasser A, Luksys G, Xin L, Pessiglione M, Gruetter R, Sandi C. Glutamine-to-glutamate ratio in the nucleus accumbens predicts effort-based motivated performance in humans. Neuropsychopharmacol. 2020;45:2048–2057.

13. Wiehler A, Branzoli F, Adanyeguh I, Mochel F, Pessiglione M. A neuro-metabolic account of why daylong cognitive work alters the control of economic decisions. Current Biology. 2022;32:3564–3575.e5.

14. Zalachoras I, Ramos-Fernández E, Hollis F, Trovo L, Rodrigues J, Strasser A, et al. Glutathione in the nucleus accumbens regulates motivation to exert reward-incentivized effort. eLife. 2022;11:e77791.

15. Gailliot MT, Baumeister RF. The Physiology of Willpower: Linking Blood Glucose to Self-Control. Pers Soc Psychol Rev. 2007;11:303–327.

16. Devine MJ, Kittler JT. Mitochondria at the neuronal presynapse in health and disease. Nat Rev Neurosci. 2018;19:63–80.

17. Lutas A, Yellen G. The ketogenic diet: metabolic influences on brain excitability and epilepsy. Trends in Neurosciences. 2013;36:32–40.

18. Morató L, Astori S, Zalachoras I, Rodrigues J, Ghosal S, Huang W, et al. eNAMPT actions through nucleus accumbens NAD+/SIRT1 link increased adiposity with sociability deficits programmed by peripuberty stress. Sci Adv. 2022;8:eabj9109.

19. Tiwari A, Myeong J, Hashemiaghdam A, Zhang H, Niu X, Laramie MA, et al. Mitochondrial pyruvate transport regulates presynaptic metabolism and neurotransmission. 2024:2024.03.20.586011.

20. Brooks GA. Lactate as a fulcrum of metabolism. Redox Biol. 2020;35:101454.

21. Gladden LB. Lactate metabolism: a new paradigm for the third millennium. J Physiol. 2004;558:5–30.

22. Dalsgaard MK. Fuelling cerebral activity in exercising man. J Cereb Blood Flow Metab. 2006;26:731–750.

23. Ide K, Secher NH. Cerebral blood flow and metabolism during exercise. Prog Neurobiol. 2000;61:397–414.

24. Quistorff B, Secher NH, Van Lieshout JJ. Lactate fuels the human brain during exercise. FASEB J. 2008;22:3443–3449.

25. Boumezbeur F, Petersen KF, Cline GW, Mason GF, Behar KL, Shulman GI, et al. The contribution of blood lactate to brain energy metabolism in humans measured by dynamic 13C nuclear magnetic resonance spectroscopy. J Neurosci. 2010;30:13983–13991.

26. van Hall G, Strømstad M, Rasmussen P, Jans O, Zaar M, Gam C, et al. Blood lactate is an important energy source for the human brain. J Cereb Blood Flow Metab. 2009;29:1121– 1129.

27. Hill AV, Long CNH, Lupton H. Muscular exercise, lactic acid and the supply and utilisation of oxygen.— Parts VII–VIII. Proceedings of the Royal Society of London Series B, Containing Papers of a Biological Character. 1924;97:155–176.

28. Hill AV, Kupalov P. Anaerobic and aerobic activity in isolated muscle. Proceedings of the Royal Society of London Series B, Containing Papers of a Biological Character. 1929;105:313–322.

29. Hirvonen J, Nummela A, Rusko H, Rehunen S, Härkönen M. Fatigue and changes of ATP, creatine phosphate, and lactate during the 400-m sprint. Canadian Journal of Sport Sciences = Journal Canadien Des Sciences Du Sport. 1992;17:141–144.

30. Tesch P, Sjödin B, Thorstensson A, Karlsson J. Muscle fatigue and its relation to lactate accumulation and LDH activity in man. Acta Physiol Scand. 1978;103:413–420.

31. Cremer JE, Braun LD, Oldendorf WH. Changes during development in transport processes of the blood-brain barrier. Biochim Biophys Acta. 1976;448:633–637.

32. Smith D, Pernet A, Hallett WA, Bingham E, Marsden PK, Amiel SA. Lactate: a preferred fuel for human brain metabolism in vivo. J Cereb Blood Flow Metab. 2003;23:658–664.

33. Magistretti PJ, Allaman I. Lactate in the brain: from metabolic end-product to signalling molecule. Nat Rev Neurosci. 2018;19:235–249.

34. Xue X, Liu B, Hu J, Bian X, Lou S. The potential mechanisms of lactate in mediating exercise-enhanced cognitive function: a dual role as an energy supply substrate and a signaling molecule. Nutr Metab (Lond). 2022;19:52.

35. Brooks GA. Cell-cell and intracellular lactate shuttles. J Physiol. 2009;587:5591–5600.

36. Mosienko V, Teschemacher AG, Kasparov S. Is L-lactate a novel signaling molecule in the brain? J Cereb Blood Flow Metab. 2015;35:1069–1075.

37. Akter M, Ma H, Hasan M, Karim A, Zhu X, Zhang L, et al. Exogenous L-lactate administration in rat hippocampus increases expression of key regulators of mitochondrial biogenesis and antioxidant defense. Front Mol Neurosci. 2023;16.

38. Descalzi G, Gao V, Steinman MQ, Suzuki A, Alberini CM. Lactate from astrocytes fuels learning-induced mRNA translation in excitatory and inhibitory neurons. Commun Biol. 2019;2:247.

39. Suzuki A, Stern SA, Bozdagi O, Huntley GW, Walker RH, Magistretti PJ, et al. Astrocyte-neuron lactate transport is required for long-term memory formation. Cell. 2011;144:810–823.

40. Dienel GA, Hertz L. Glucose and lactate metabolism during brain activation. J Neurosci Res. 2001;66:824–838.

41. Hladky SB, Barrand MA. Metabolite Clearance During Wakefulness and Sleep. Handb Exp Pharmacol. 2019;253:385–423.

42. Lundgaard I, Lu ML, Yang E, Peng W, Mestre H, Hitomi E, et al. Glymphatic clearance controls state-dependent changes in brain lactate concentration. J Cereb Blood Flow Metab. 2017;37:2112–2124.

43. Theriault JE, Shaffer C, Dienel GA, Sander CY, Hooker JM, Dickerson BC, et al. A functional account of stimulation-based aerobic glycolysis and its role in interpreting BOLD signal intensity increases in neuroimaging experiments. Neurosci Biobehav Rev. 2023;153:105373.

44. Kahneman D, Tversky A. Choices, values, and frames. American Psychologist. 1984;39:341– 350.

45. Hayes DJ, Northoff G. Common brain activations for painful and non-painful aversive stimuli. BMC Neuroscience. 2012;13:60.

46. Bartra O, McGuire JT, Kable JW. The valuation system: A coordinate-based meta-analysis of BOLD fMRI experiments examining neural correlates of subjective value. NeuroImage. 2013;76:412–427.

47. Knight FH, Frank H. Risk, uncertainty and profit. Boston, New York, Houghton Mifflin Company; 1921.

48. Lopez-Gamundi P, Mas-Herrero E, Marco-Pallares J. Disentangling effort from probability of success: Temporal dynamics of frontal midline theta in effort-based reward processing. Cortex. 2024;176:94–112.

49. Lee CY, Soliman H, Geraghty BJ, Chen AP, Connelly KA, Endre R, et al. Lactate topography of the human brain using hyperpolarized 13C-MRI. Neuroimage. 2020;204:116202.

50. Kurniawan IT, Grueschow M, Ruff CC. Anticipatory Energization Revealed by Pupil and Brain Activity Guides Human Effort-Based Decision Making. J Neurosci. 2021;41:6328–6342.

51. Touroutoglou A, Andreano J, Dickerson BC, Barrett LF. The tenacious brain: How the anterior mid-cingulate contributes to achieving goals. Cortex. 2020;123:12–29.

52. Müller T, Klein-Flügge MC, Manohar SG, Husain M, Apps MAJ. Neural and computational mechanisms of momentary fatigue and persistence in effort-based choice. Nat Commun. 2021;12:4593.

53. Lim S-I, Xin L. γ-aminobutyric acid measurement in the human brain at 7 T: Short echo-time or Mescher–Garwood editing. NMR in Biomedicine. 2022;35:e4706.

54. Cuenoud B, Huang Z, Hartweg M, Widmaier M, Lim S, Wenz D, et al. Effect of circadian rhythm on NAD and other metabolites in human brain. Front Physiol. 2023;14:1285776.

55. Soutschek A, Nadporozhskaia L, Christian P. Brain stimulation over dorsomedial prefrontal cortex modulates effort-based decision making. Cogn Affect Behav Neurosci. 2022;22:1264– 1274.

56. Chong TT-J, Apps M, Giehl K, Sillence A, Grima LL, Husain M. Neurocomputational mechanisms underlying subjective valuation of effort costs. PLoS Biol. 2017;15:e1002598.

57. Le Bouc R, Pessiglione M. A neuro-computational account of procrastination behavior. Nat Commun. 2022;13:5639.

58. Yao Y-W, Song K-R, Schuck NW, Li X, Fang X-Y, Zhang J-T, et al. The dorsomedial prefrontal cortex represents subjective value across effort-based and risky decision-making. Neuroimage. 2023;279:120326.

59. Clairis N, Pessiglione M. Value, confidence, deliberation: a functional partition of the medial prefrontal cortex demonstrated across rating and choice tasks. J Neurosci. 2022;42:5580– 5592.

60. Grinband J, Savitskaya J, Wager TD, Teichert T, Ferrera VP, Hirsch J. The dorsal medial frontal cortex is sensitive to time on task, not response conflict or error likelihood. Neuroimage. 2011;57:303–311.

61. McGuire JT, Botvinick MM. Prefrontal cortex, cognitive control, and the registration of decision costs. Proc Natl Acad Sci U S A. 2010;107:7922–7926.

62. Prévost C, Pessiglione M, Météreau E, Cléry-Melin M-L, Dreher J-C. Separate valuation subsystems for delay and effort decision costs. J Neurosci. 2010;30:14080–14090.

63. Arulpragasam AR, Cooper JA, Nuutinen MR, Treadway MT. Corticoinsular circuits encode subjective value expectation and violation for effortful goal-directed behavior. Proc Natl Acad Sci U S A. 2018;115:E5233–E5242.

64. Clairis N, Lopez-Persem A. Debates on the dorsomedial prefrontal/dorsal anterior cingulate cortex: insights for future research. Brain. 2023:awad263.

65. Clairis N, Pessiglione M. Value estimation versus effort mobilization: a general dissociation between ventromedial and dorsomedial prefrontal cortex. J Neurosci. 2024. 20 March 2024. 10.1523/JNEUROSCI.1176-23.2024.

66. Skvortsova V, Palminteri S, Pessiglione M. Learning To Minimize Efforts versus Maximizing Rewards: Computational Principles and Neural Correlates. Journal of Neuroscience. 2014;34:15621–15630.

67. André N, Audiffren M, Baumeister RF. An Integrative Model of Effortful Control. Front Syst Neurosci. 2019;13:79.

68. Feng C, Thompson WK, Paulus MP. Effect sizes of associations between neuroimaging measures and affective symptoms: A meta-analysis. Depression and Anxiety. 2022;39:19–25.

69. Marek S, Tervo-Clemmens B, Calabro FJ, Montez DF, Kay BP, Hatoum AS, et al. Reproducible brain-wide association studies require thousands of individuals. Nature. 2022;603:654–660.

70. Allen D, Westerblad H. Lactic Acid--The Latest Performance-Enhancing Drug. Science. 2004;305:1112–1113.

71. Cai Y, Guo H, Han T, Wang H. Lactate: a prospective target for therapeutic intervention in psychiatric disease. Neural Regen Res. 2024;19:1473–1479.

72. Carrard A, Elsayed M, Margineanu M, Boury-Jamot B, Fragnière L, Meylan EM, et al. Peripheral administration of lactate produces antidepressant-like effects. Mol Psychiatry. 2018;23:392–399.

73. Karnib N, El-Ghandour R, El Hayek L, Nasrallah P, Khalifeh M, Barmo N, et al. Lactate is an antidepressant that mediates resilience to stress by modulating the hippocampal levels and activity of histone deacetylases. Neuropsychopharmacology. 2019;44:1152–1162.

74. Cauli B, Dusart I, Li D. Lactate as a determinant of neuronal excitability, neuroenergetics and beyond. Neurobiol Dis. 2023;184:106207.

75. Karagiannis A, Gallopin T, Lacroix A, Plaisier F, Piquet J, Geoffroy H, et al. Lactate is an energy substrate for rodent cortical neurons and enhances their firing activity. Elife. 2021;10:e71424.

76. Yao S, Xu M-D, Wang Y, Zhao S-T, Wang J, Chen G-F, et al. Astrocytic lactate dehydrogenase A regulates neuronal excitability and depressive-like behaviors through lactate homeostasis in mice. Nat Commun. 2023;14:729.

77. Le Bouc R, Borderies N, Carle G, Robriquet C, Vinckier F, Daunizeau J, et al. Effort avoidance as a core mechanism of apathy in frontotemporal dementia. Brain. 2022:awac427.

78. Trevisi G, Eickhoff SB, Chowdhury F, Jha A, Rodionov R, Nowell M, et al. Probabilistic electrical stimulation mapping of human medial frontal cortex. Cortex. 2018;109:336–346.

79. Cremer JE, Cunningham VJ, Pardridge WM, Braun LD, Oldendorf WH. Kinetics of blood-brain barrier transport of pyruvate, lactate and glucose in suckling, weanling and adult rats. J Neurochem. 1979;33:439–445.

80. Hladky SB, Barrand MA. Fluid and ion transfer across the blood-brain and blood-cerebrospinal fluid barriers; a comparative account of mechanisms and roles. Fluids Barriers CNS. 2016;13:19.

81. Todd JJ. Lactate: valuable for physical performance and maintenance of brain function during exercise. Bioscience Horizons: The International Journal of Student Research. 2014;7:hzu001.

82. Ishii H, Nishida Y. Effect of Lactate Accumulation during Exercise-induced Muscle Fatigue on the Sensorimotor Cortex. J Phys Ther Sci. 2013;25:1637–1642.

83. Bianchi MC, Sgandurra G, Tosetti M, Battini R, Cioni G. Brain magnetic resonance in the diagnostic evaluation of mitochondrial encephalopathies. Biosci Rep. 2007;27:69–85.

84. Maddock RJ, Buonocore MH. MR spectroscopic studies of the brain in psychiatric disorders. Curr Top Behav Neurosci. 2012;11:199–251.

85. Mascalchi M, Montomoli M, Guerrini R. Neuroimaging in mitochondrial disorders. Essays Biochem. 2018;62:409–421.

86. Li J, Xia Y, Xu H, Xiong R, Zhao Y, Li P, et al. Activation of brain lactate receptor GPR81 aggravates exercise-induced central fatigue. Am J Physiol Regul Integr Comp Physiol. 2022;323:R822–R831.

87. Demello JJ, Cureton KJ, Boineau RE, Singh MM. Ratings of perceived exertion at the lactate threshold in trained and untrained men and women. Med Sci Sports Exerc. 1987;19:354–362.

88. Hetzler RK, Seip RL, Boutcher SH, Pierce E, Snead D, Weltman A. Effect of exercise modality on ratings of perceived exertion at various lactate concentrations. Med Sci Sports Exerc. 1991;23:88–92.

89. Scherr J, Wolfarth B, Christle JW, Pressler A, Wagenpfeil S, Halle M. Associations between Borg’s rating of perceived exertion and physiological measures of exercise intensity. Eur J Appl Physiol. 2013;113:147–155.

90. Béland-Millar A, Larcher J, Courtemanche J, Yuan T, Messier C. Effects of Systemic Metabolic Fuels on Glucose and Lactate Levels in the Brain Extracellular Compartment of the Mouse. Front Neurosci. 2017;11:7.

91. Murack M, Messier C. The impact of lactic acid and medium chain triglyceride on blood glucose, lactate and diurnal motor activity: A re-examination of a treatment of major depression using lactic acid. Physiology & Behavior. 2019;208:112569.

92. Ellis D, Simmons C, Miller BF. Sodium lactate infusion during a cycling time-trial does not increase lactate concentration or decrease performance. European Journal of Sport Science. 2009;9:367–374.

93. Pollak KA, Swenson JD, Vanhaitsma TA, Hughen RW, Jo D, White AT, et al. Exogenously applied muscle metabolites synergistically evoke sensations of muscle fatigue and pain in human subjects. Exp Physiol. 2014;99:368–380.

94. Rae CD, Baur JA, Borges K, Dienel G, Díaz-García CM, Douglass SR, et al. Brain energy metabolism: A roadmap for future research. J Neurochem. 2024. 6 January 2024. 10.1111/jnc.16032.

95. Veech RL. The metabolism of lactate. NMR Biomed. 1991;4:53–58.

96. Morant-Ferrando B, Jimenez-Blasco D, Alonso-Batan P, Agulla J, Lapresa R, Garcia-Rodriguez D, et al. Fatty acid oxidation organizes mitochondrial supercomplexes to sustain astrocytic ROS and cognition. Nat Metab. 2023;5:1290–1302.

97. York EM, Miller A, Stopka SA, Martínez-François JR, Hossain MA, Baquer G, et al. The dentate gyrus differentially metabolizes glucose and alternative fuels during rest and stimulation. J Neurochem. 2024;168:533–554.

98. Dembitskaya Y, Piette C, Perez S, Berry H, Magistretti PJ, Venance L. Lactate supply overtakes glucose when neural computational and cognitive loads scale up. Proceedings of the National Academy of Sciences. 2022;119:e2212004119.

99. Chow LS, Gerszten RE, Taylor JM, Pedersen BK, van Praag H, Trappe S, et al. Exerkines in health, resilience and disease. Nat Rev Endocrinol. 2022;18:273–289.

100. Brooks GA, Osmond AD, Arevalo JA, Curl CC, Duong JJ, Horning MA, et al. Lactate as a major myokine and exerkine. Nat Rev Endocrinol. 2022;18:712.

101. Chow LS, Gerszten RE, Taylor JM, Pedersen BK, van Praag H, Trappe S, et al. Reply to ‘Lactate as a major myokine and exerkine’. Nat Rev Endocrinol. 2022;18:713–713.

102. Chen Y, Lin Q, Liao X, Zhou C, He Y. Association of aerobic glycolysis with the structural connectome reveals a benefit-risk balancing mechanism in the human brain. Proc Natl Acad Sci U S A. 2021;118:e2013232118.

103. Goyal MS, Hawrylycz M, Miller JA, Snyder AZ, Raichle ME. Aerobic glycolysis in the human brain is associated with development and neotenous gene expression. Cell Metab. 2014;19:49–57.

104. Vaishnavi SN, Vlassenko AG, Rundle MM, Snyder AZ, Mintun MA, Raichle ME. Regional aerobic glycolysis in the human brain. Proc Natl Acad Sci U S A. 2010;107:17757–17762.

105. Castrillon G, Epp S, Bose A, Fraticelli L, Hechler A, Belenya R, et al. An energy costly architecture of neuromodulators for human brain evolution and cognition. Science Advances. 2023;9:eadi7632.

106. Blain B, Hollard G, Pessiglione M. Neural mechanisms underlying the impact of daylong cognitive work on economic decisions. Proceedings of the National Academy of Sciences. 2016;113:6967–6972.

107. Schmidt L, Lebreton M, Cléry-Melin M-L, Daunizeau J, Pessiglione M. Neural Mechanisms Underlying Motivation of Mental Versus Physical Effort. PLoS Biology. 2012;10:e1001266.

108. Soutschek A, Tobler PN. Causal role of lateral prefrontal cortex in mental effort and fatigue. Hum Brain Mapp. 2020;41:4630–4640.

109. Suzuki S, Lawlor VM, Cooper JA, Arulpragasam AR, Treadway MT. Distinct regions of the striatum underlying effort, movement initiation and effort discounting. Nat Hum Behav. 2021;5:378–388.

110. Brown JW, Alexander WH. Foraging Value, Risk Avoidance, and Multiple Control Signals: How the Anterior Cingulate Cortex Controls Value-based Decision-making. Journal of Cognitive Neuroscience. 2017;29:1656–1673.

111. Shenhav A, Botvinick MM, Cohen JD. The expected value of control: an integrative theory of anterior cingulate cortex function. Neuron. 2013;79:217–240.

112. Barakat A, Clairis N, Brochard J, Xin L, Sandi C. Predicting individual variations in mental effort-based decision-making using machine learning: Neurometabolic signature in the dorsomedial prefrontal cortex/dorsal anterior cingulate cortex. 2024:2024.01.23.576854.

113. Bondolfi G, Jermann F, Rouget BW, Gex-Fabry M, McQuillan A, Dupont-Willemin A, et al. Self- and clinician-rated Montgomery-Asberg Depression Rating Scale: evaluation in clinical practice. J Affect Disord. 2010;121:268–272.

114. Yee A, Yassim ARM, Loh HS, Ng CG, Tan K-A. Psychometric evaluation of the Malay version of the Montgomery-Asberg Depression Rating Scale (MADRS-BM). BMC Psychiatry. 2015;15:200.

115. Brainard DH. The Psychophysics Toolbox. Spatial Vision. 1997;10:433–436.

116. Daunizeau J, Adam V, Rigoux L. VBA: A Probabilistic Treatment of Nonlinear Models for Neurobiological and Behavioural Data. PLoS Comput Biol. 2014;10:e1003441.

117. Diedenhofen B, Musch J. cocor: A Comprehensive Solution for the Statistical Comparison of Correlations. PLOS ONE. 2015;10:e0121945.

118. Steiger JH. Tests for comparing elements of a correlation matrix. Psychological Bulletin. 1980;87:245–251.

119. Dei Cas M, Paroni R, Saccardo A, Casagni E, Arnoldi S, Gambaro V, et al. A straightforward LC-MS/MS analysis to study serum profile of short and medium chain fatty acids. Journal of Chromatography B. 2020;1154:121982.

120. Gruetter R. Automatic, localized in vivo adjustment of all first- and second-order shim coils. Magn Reson Med. 1993;29:804–811.

121. Marques JP, Kober T, Krueger G, van der Zwaag W, Van de Moortele P-F, Gruetter R. MP2RAGE, a self bias-field corrected sequence for improved segmentation and T1-mapping at high field. NeuroImage. 2010;49:1271–1281.

122. Afif A, Mertens P. Description of sulcal organization of the insular cortex. Surg Radiol Anat. 2010;32:491–498.

123. Mekle R, Mlynárik V, Gambarota G, Hergt M, Krueger G, Gruetter R. MR spectroscopy of the human brain with enhanced signal intensity at ultrashort echo times on a clinical platform at 3T and 7T. Magn Reson Med. 2009;61:1279–1285.

124. Öz G, Deelchand DK, Wijnen JP, Mlynárik V, Xin L, Mekle R, et al. Advanced single voxel 1 H magnetic resonance spectroscopy techniques in humans: Experts’ consensus recommendations. NMR Biomed. 2020:e4236.

125. Xin L, Schaller B, Mlynarik V, Lu H, Gruetter R. Proton T1 relaxation times of metabolites in human occipital white and gray matter at 7 T. Magn Reson Med. 2013;69:931–936.

126. Provencher SW. Estimation of metabolite concentrations from localizedin vivo proton NMR spectra. Magn Reson Med. 1993;30:672–679.

127. Kriegeskorte N, Simmons WK, Bellgowan PSF, Baker CI. Circular analysis in systems neuroscience: the dangers of double dipping. Nat Neurosci. 2009;12:535–540.

128. Rosseel Y. lavaan: An R Package for Structural Equation Modeling. Journal of Statistical Software. 2012;48:1–36.

