## Supplementary material for "A neurometabolic mechanism involving dmPFC/dACC lactate in physical effort-based decision-making"

### Supplementary Methods

#### Online Questionnaires

Before coming to the laboratory for the experiment, participants completed several online questionnaires. These surveys aimed at establishing a better profile of the participants and also to ensure a wide range of motivational profiles in our study (see *Materials & Methods*). The questionnaires focused on assessing different personality dimensions including motivation, stress and anxiety, early-life stress, dominance and competitiveness, as well as general metabolic and psychological characteristics (socio-economic status, weight, height and HEXACO for the general profile). The questionnaires related to stress and anxiety were the following: French version of the Perceived Stress Scale (PSS-14) [1], the French version of the trait part of the Spielberger State-Trait Anxiety Inventory (STAI-T) [2] and a french translation of the Social Interaction Anxiety Scale (SIAS) [3].

#### Subjective Stress and Fatigue Ratings

Right before going in the scanner for ^1^H-MRS or fMRI measurement and just after getting out of the scanner, participants were required to provide a rating of their current subjective level of fatigue and of stress on a visual analogue scale going from 0 to 10 to monitor their levels of stress and fatigue across the duration of the experiment.

#### Sleep Questionnaire

At the same moment they filled the stress and fatigue questionnaires, participants were also required to indicate their average amount of sleep (in hours and minutes) and the approximate amount of sleep they had the day before the experiment (in hours and minutes).

#### Saliva collection

Every time they filled the stress, fatigue and sleep questionnaires, we also collected participants’ saliva using 2.0 mL passive-drool collection tubes (Salimetrics, USA). The tubes were immediately put in dry ice after saliva collection and then they were stored in a -80°C freezer. We therefore collected four salivary tubes per participant, one before and one after each session in the MRI scanner, aiming at measuring salivary testosterone and cortisol levels.

#### Salivary Cortisol Analysis

Saliva samples were centrifuged at 3000 rpm for 15 min at 4°C and they were then quantified with the Salimetrics Salivary Cortisol Enzyme linked immunosorbent assay (ELISA) kit according to the manufacturer’s instructions (Salimetrics, USA).

#### Behavioral Task

*Training and calibration.* After ^1^H-MRS and before performing the actual task in fMRI, participants were trained for a period lasting 44±1 minutes (mean ± sem). To ensure proper calibration, participants were instructed that during a measure of their maximal physical and mental capacity, they could win up to 4 CHF based on their performance each time. During physical calibration, participants had to squeeze a dynamometer (TSD121B-MRI, BIOPAC Inc.) as hard as possible using their left hand within a 5 seconds period. Calibration was repeated three times. The maximum of these three attempts was defined as their maximal voluntary contraction (MVC). During a physical effort, participants had to exert a force at least higher than 55% of their MVC. During mental calibration, participants were extensively trained to perform a 2-back task as fast as possible until they could achieve 6 correct responses under 10 seconds, at least 80% of the time. Following this learning period, they were instructed to achieve the highest possible score in under 10 seconds, with 6 responses being the minimum requirement which served as a calibration for the difficulty levels. After three attempts, we defined 80% of their averaged best score as the maximum number of correct responses (MNCR), in order to consider the high variability in this measure and to minimize the confusion between risk and effort aversion. In the physical effort task, the difficulty varied in terms of duration that someone had to squeeze the handgrip above 55% of their MVC (0.5/1.5/2.5/4.5s), while in the mental effort task, the difficulty varied in terms of number of correct answers $(\frac{1}{9}/\frac{1}{3}/\frac{2}{3}/1)*MNCR$ to provide within 10s.

For each effort type (physical/mental) calibration was followed by a training period where participants had to perform all four effort levels 5 times each in order to get familiarized with the range of difficulty expected in the main task. After this effort training, participants received a short training for the actual task consisting in 4 choice trials, presented as in the main task, followed by effort performance to ensure that they understood the way the task would work.

*Indifference point measurement.* At the end of the training session for each effort type, we calibrated the incentive levels by computing an indifference point (IP) independently for each effort type following a short IP measurement procedure adapted from Westbrook and colleagues [4, 5]. This procedure consisted in presenting 5 subsequent choice trials and asking to the participant their subjective preference on each trial. After their choice was selected, they always had to perform the associated effort subsequently and their earnings in this period would be added to their final monetary outcome, so that their choices were consequential. Choices were always between a small and fixed reward (0.5 CHF) for the easiest effort level (effort level 0) and a larger, varying reward for the medium level of effort (effort level 2). The large reward was of 1.00 CHF on the first trial and, after each choice, it either increased or decreased depending on whether the subject picked the low or the high effort option, respectively. The change in value for the high reward varied following $\frac{hR-0.5}{2}$ with $hR$ representing the momentary monetary amount of the high effort option on each trial, such that the incentive value converged to the participant’s indifference point (IP) over 5 trials. For the loss condition and for the sake of time, the indifference point was defined by symmetry as -0.5-Δ. For each condition (gain/loss) and each effort type (physical/mental), the three incentive levels proposed were computed as $IP-\frac{\Delta R}{2}$, $IP$, and $IP+\frac{\Delta R}{2}$ where $\Delta R=IP-0.5$. At the end of the IP measurement, participants went back in the MRI scanner to perform the behavioral task during fMRI.

#### Behavioral Analysis

All data were analyzed using MATLAB 2021a (The MathWorks Inc.).

To extract motivational parameters from participants’ behavior, choices were fitted using Matlab VBA toolbox (<https://mbb-team.github.io/VBA-toolbox/>) which implements Variational Bayesian analysis under the Laplace approximation [6]. The algorithm provides an estimate of the posterior density over the model-free parameters, starting with Gaussian priors. Uninformed priors (N(0, 100)) were used to fit the model on participants’ behavior.

Our model aimed at characterizing the probability of choosing the high effort option. Sessions where participants systematically picked up either the high or the low effort option in 100% of the trials were ignored from the analysis. To further inform our models, we included confidence levels for each choice so that there were four possible choices on each trial: low effort with high confidence (0), low effort with low confidence (0.25), high effort with low confidence (0.75), or high effort with high confidence (1.00). In each trial *t*, our model assumes that individuals compare the subjective values (SV) of the two options and select the best option accordingly:

$\Delta SV\left( t \right)= kR\cdot\Delta R\left( t \right)+kP\cdot\Delta P(t)-\Delta E_{p}(t)*\left( kE_{p}+kF_{p}\cdot Fp(t) \right))-\Delta E_{m}(t)*\left( kE_{m}-kL_{m}\cdot ME(t) \right)$ (Eq. 1)

Where $t$ is the trial number, ∆SV is the difference in subjective value (SV) between the low and the high effort options, ΔR is the difference in reward amounts between the high and the low effort options (equal to 0 in punishment trials), ΔP is the difference in punishment amounts between the high and the low effort options (equal to 0 in reward trials), ΔEp is the difference in physical effort levels between the high and the low effort options (equal to 0 in mental effort blocks), ΔEm is the difference in mental effort levels between the high and the low effort options (equal to 0 in physical effort blocks), Fp represents the accumulated force exerted in the physical task and ME the momentary efficiency in performing the mental effort task. The model allows to estimate 7 behavioral parameters which consist in the sensitivity to reward kR, the sensitivity to punishments kP, the sensitivity to physical effort kEp, the sensitivity to mental effort kEm, the sensitivity to physical fatigue kFp and the sensitivity to mental learning kLm. The fatigue component in the physical task represents the sum of exerted force until trial $t$ and is defined according to the formula $F_{p}(t)= \sum_{i=0}^{t-1} AUC(i)$ with AUC corresponding to the area under the curve of the force exerted in trial $i$. In the mental task, we included a momentary efficiency component $ME(t)=\frac{Nb correct answers(t-1)}{Total trial time(t-1)}$ which represents the momentary task performance which seemed to improve over time, possibly due to learning. Finally, the choice probability is modeled by a *softmax* function including a bias constant (kBias):

$P_{HE} \left( t \right)=\frac{1}{1+e^{-\Delta SV(t)+kBias}}$ (Eq. 2)

Where $P_{HE}\left( t \right)$represents the probability of choosing the high effort (HE) option. Apart from the bias parameter (*kBias*), all parameters were positively constrained with a ${log}^{1+e(x)}$ transformation. Priors were quasi-uniform as defined by a centered gaussian with large standard deviation (100), accounting for high fluctuations in ∆R and ∆P across participants.

To assess the model’s validity, we performed simulations in order to verify parameters identifiability and recoverability [7]. We launched 10 000 simulations based on the average ΔR, ΔP, ΔEp, and ΔEm across subjects and by using the average AUC and ME for each effort level across subjects. Fp and ME were updated on each trial based on the choice probability output from the model. We simulated 10 000 sets of normally distributed parameters centered on 0 and then tested the recoverability and identifiability of these parameters using our behavioral model.

To further challenge our model’s assumptions, we ran a simplified frequentist version of it, i.e. without gaussian priors or positivity constraints and replacing the physical fatigue and mental learning by trial number to verify that time had an impact on HE choices and, if so, whether the direction was the same as the one imposed in our model. Choices were modeled separately with a logistic regression in each task following the formula:

$P_{HE} \left( t \right)= \frac{1}{1+e^{-(\beta R\cdot\Delta R(t)+\beta P\cdot\Delta P(t) -(\beta E+ \beta T\cdot t)\cdot\Delta E(t)+\beta Bias)+\varepsilon}}$ (Eq. 3)

In this equation, $\beta R$ and $\beta P$ represent the sensitivity to monetary incentives (reward and punishment, respectively), $\beta E$ represents the aversion to effort, $\beta T$ represents the influence of time on the decision and $\beta Bias$ is a constant corresponding to a general bias for the high or the low effort option, independent of the other variables. $\Delta R$ and $\Delta P$ corresponded to the difference between the reward (or punishment respectively) amount associated to the high *versus* low effort options, which were established individually based on the indifference point calibration. $\Delta E$ corresponded to the difference between the effort level associated to the high effort option (1/2/3) and the effort associated to the low effort options (0). $t$ corresponds to the trial number within the session (1 to 54) and was present in order to take the effect of time on choices into consideration, and ε is the error term. The choices were fitted independently for the physical (p) and the mental (m) effort task and for each subject with this model. Then, the betas were tested at the group-level with a t.test against zero.

To explore the potential link between choice deliberation times (DT) and the task variables, we decided to model DT according to monetary incentives, effort levels, trial valence and confidence given that DT are known to vary with reward, effort and confidence [8, 9]. DT were defined as the time between stimulus onset and the first button press [10]. Trial-wise variations in DT were fitted with a linear regression model. This model, including a session-specific intercept (β_s1_/β_s2_/β_s3_/β_s4_) and a noise term ε, was designed as follows:

$DT = {\varepsilon+\beta}_{s1}+\beta_{s2}+\beta_{s3}+\beta_{s4}+\beta_{R}\cdot\Delta R+\beta_{P}\cdot\Delta P+\beta_{RP}\cdot RP+\beta_{Ep}\cdot\Delta Ep+\beta_{Em}\cdot\Delta Em+\beta_{Conf}\cdot Conf+\beta_{t}\cdot t$ (Eq. 4)

In this equation, $Conf$ was defined as $\boldsymbol{(}P_{HE}-0.5\boldsymbol{)^{2}}$ based on previous research which shows that this is a good proxy for subjective confidence [10] which is more gradual than the binary confidence judgments provided by the participants. The other variables were similar to the model above (**Eq. 3**), with $t$ representing trial number to account for the effect of time on choice DT and RP being a trial-type variable encoding reward trials (as ones) and punishment trials (as zeros) which effectively captures a general DT bias depending on trial valence with appetitive choice trials being generally faster than more aversive trials.

We also controlled for the correlation between our main variables of interest (Ech, SVch, DT) by performing 2-by-2 correlations at the individual level to extract individual correlation coefficients r and R² parameters, after zscoring each variable within each block, as was done for the fMRI regressors. We then tested whether the correlation coefficients r were significantly different from zero at the group-level by using a Student’s t.test against zero. We first performed these correlation tests across all tasks and then we extracted the variables within each task to see if the results were similar or not.

#### fMRI data analysis

Given that the subjective value of the chosen option (SVch), the effort chosen (Ech) and choice deliberation times (DT) might not be completely orthogonal to each other, we decided to build two further GLMs, GLM2 and GLM3, to further assess whether the results observed in GLM1 were holding independent of the way the regressors were orthogonalized to each other. GLM2 and GLM3 were completely equivalent to GLM1, but, in GLM2, the regressors were orthogonalized in the DT/SVch/Ech order to account for DT and SVch influence on Ech, while, in GLM3, the regressors were orthogonalized in the Ech/SVch/DT order to account for the influence of Ech on SVch and DT.

Moreover, we wanted to control that the correlations we observed between dmPFC/dACC activity, aIns activity and Ech were not due to the dmPFC/dACC and/or aIns activity correlating with the effort costs of the HE option, which could mean that dmPFC/dACC and/or aIns are not promoting HE selection, but are instead encoding the costs of the HE option, therefore potentially informing other areas performing the decision. For this purpose, we designed GLM4 which included the same categorical regressors as GLM1, except for the choice period regressor. The choice period was modeled as two categorical regressors depending on the choice made: one for the trials where the low effort was chosen (lEch) and another for the trials where the high effort was chosen (hEch). Both regressors were parametrically modulated by the level of the high effort (HE) option, the subjective value of the high effort option (hSV) and the deliberation time (DT). Because of the low variability in participants’ choices in some blocks (for example, some subject would pick the low effort option only when the HE level was 3, meaning no variability for the HE parametric modulator of the lEch regressor), we had to remove a few more blocks and/or subjects from this analysis leading to a final N = 60 subjects included.

### Supplementary Results

#### Validating The Task Parameters

Model recoverability was evidenced by high correlations between true and estimated parameters (r ≥ 0.79). Identifiability was supported by low inter-parameter correlations for the estimated parameters (|r| < 0.10). Moreover, the model accurately mirrored participants’ choices (median absolute error (MAE) of 20.5 ± 0.8%, goodness of fit R² = 0.46).

To further validate the results of our computational model, we used a conventional generalized linear model (GLM) with no constraint on the parameters. Based on a basic logistic regression (Eq. 4), we observed that high effort (HE) options were indeed selected significantly more often when participants were offered higher rewards ($\beta R_{m}$ = 50.13 ± 7.09, mean ± sem; p < 0.001), lower punishments ($\beta P_{m}$ = 30.95 ± 4.04; p < 0.001) and lower physical ($\beta E_{p}$= 1.65 ± 0.14; p < 0.001) or mental ($\beta E_{m}$= 1.94 ± 0.14; p < 0.001) effort amounts, confirming the validity of the parameters used in our model and of the positivity constraint we forced upon them to increase model’s reliability. Moreover, high physical effort (HPE) options were selected significantly less often across trials ($\beta T_{p}$= 1.6·10^-4^ ± 2.2·10^-5^; p < 0.001) validating that participants were subject to physical fatigue (or at least that they actively tried to prevent its emergence). On the contrary, in the mental effort task, they chose more often the high mental effort (HME) option across trials ($\beta T_{m}$ = -0.21 ± 0.04; p < 0.001) possibly because, despite the extensive preceding training, their efficiency in doing the mental efforts improved over time (β = 0.010 ± 0.003; p = 0.003). Interestingly, participants also displayed a significant bias towards selecting the HE option independent of the varying variables of the task (βBias = 2.58 ± 0.23; p < 0.001), further confirming the validity of the parameters included in our main computational model.

We initially expected that participants might be more motivated to choose high efforts in the loss domain than in the gain domain due to loss aversion. Quite expectedly, we observed that the proportion of HE choices in reward trials was highly correlated with the proportion of HE choices in the loss trials (r = 0.47, p < 0.001) which was also reflected in the correlation between the gain and loss parameters from our computational model (r = 0.45, p < 0.001). However, instead of loss aversion, we actually observed that the proportion of HE choices was higher in the gain than in the loss condition (gains: 69.21 ± 1.84% > losses: 61.98 ± 1.97%; p < 0.001). The same was true when looking separately at physical efforts (gains: 63.26 ± 2.55% > losses: 53.24 ± 2.76%; p < 0.001) and the same tendency appeared for mental efforts albeit not significant (gains: 76.73 ± 2.18% > losses: 73.25 ± 2.28%; p = 0.059). Accordingly, we observed that the gain parameter from our computational model also tended to be higher than the loss parameter, although not significantly (paired t.test: kR = 0.53 ± 0.12 > kP = 0.32 ± 0.06; p = 0.056). Our results therefore suggest that, in our tasks, individuals do not display loss aversion, but instead a slight boost towards exerting more effort for gains in comparison with losses.

#### ****Deliberation Times Analysis****

Next, a GLM was employed to analyze deliberation times (DT) during choices, incorporating variables such as incentive levels, effort difficulty, a bias towards reward versus punishment, and confidence levels. Participants took faster decisions (i.e., faster DTs) for rewards compared to punishments (β_RP_ = -0.189 ± 0.037; p < 0.001), but different reward levels (β_R_ = -0.917 ± 0.641; p = 0.158) or punishment levels did not significantly affect DTs (β_P_ = 1.134 ± 0.595; p = 0.061). Increased effort difficulty led to longer DTs in both the physical (Fig. S1-1B; β_Ep_ = 0.043 ± 0.015; p = 0.005) and the mental (Fig. S1-1B; β_Em_ = 0.078 ± 0.021; p < 0.001) tasks. Higher confidence levels corresponded to faster DTs (β_Conf_ = -0.464 ± 0.050; p < 0.001) as usually observed in decision-making tasks [10, 11]. DTs also decreased over successive trials (β_t_ = -0.005 ± 0.001; p < 0.001), suggesting more efficient decision-making over time. These findings align with established models of response speed and vigor, highlighting the influence of perceived value [8, 12] and task familiarity [13] on the speed of decision-making.

#### Controlling For The Relationship Between Lactate, Physical Capacity, Sex, Sleep And Stress

Because lactate was found to influence physical effort sensitivity, we decided to control that this relationship could not be explained by other factors than motivation and effort perception. More specifically, we wondered whether lactate levels would relate to differences in physical capacity, in sex (as men are generally stronger than women), in sleep, or in stress- and anxiety-related measures.

Concerning physical capacity, we found no significant correlation between plasma, dmPFC/dACC, or aIns lactate levels and maximum voluntary contraction (MVC) force (Fig. S2-1A, all p > 0.4), suggesting that lactate levels were not related to inter-individual differences in the physical capacity for effort. Because exercise-derived metabolites are often associated with improvements in cognition [14], we also assessed whether there was any relationship between our lactate measurements and the maximum number of correct responses (MNCR) performed during the mental calibration and could observe that there was no significant relationship between any of the lactate measures and MNCR (Fig. S2-1B, all p > 0.3). Furthermore, lactate levels in plasma or across the measured brain regions showed no correlation with subjective fatigue ratings (Fig. S2-1C), collected before and after MRS and fMRI sessions (all p > 0.2), or the difference in fatigue between the end and the start of the experiment (all p > 0.15), or the difference between the end and the start of the task (all p > 0.3). There was also no correlation between any of the lactate measures (plasma, dmPFC/dACC, aIns) and the difference in maximal performance before and after each fMRI block neither in the mental (all p > 0.2) nor in the physical (all p > 0.1) effort tasks (Fig. S2-1D), further suggesting the absence of relationship between our baseline lactate measures and physical or mental exertion capacities or fatigue-related measures in the task.

We also verified that there was no significant influence of sex differences on behavioral measures of motivation (Fig. S2-2A), including the proportion of HPE choices (p = 0.468) or kEp (p = 0.241), or on dmPFC/dACC (p = 0.874), aIns (p = 0.146) or plasma (p = 0.819) lactate (Fig. S2-2B) measures, further confirming that the effect of lactate we observed was not related to sex differences or to maximal capacity differences between our individuals.

Because lactate is cleared by the glymphatic system during sleep [15, 16], we also explored if there was any relationship between the average amount of sleep, the amount of sleep the day before the experiment, or the delta between the amount of sleep the day before the experiment and the average amount of sleep of each individual and plasma levels in the plasma, the dmPFC/dACC and the aIns. There was no significant relationship with either plasma (all p > 0.5), the dmPFC/dACC (all p > 0.2) or the aIns (all p > 0.3) lactate levels and any of the sleep-related measures (Fig. S2-3).

Additionally, because lactate may be altered by stress in differential manners depending on the brain area [17], we also controlled that there was no significant correlation between lactate measures and anxiety or stress-related variables. We found that there was no significant correlation between lactate levels in the brain or plasma and stress-related questionnaires (Fig. S2-4A-C, all p > 0.2), stress ratings on the day of the experiment (Fig. S2-4D, all p > 0.1), or cortisol measurements, which decreased with time-on-task according to circadian rhythm (Fig. S2-4E), on the day of the experiment (Fig. S2-4F, all p > 0.1).There was only one small negative correlation between plasma lactate and the difference between final and initial levels of cortisol (r = -0.311; p = 0.019) suggesting that higher initial plasma lactate levels correlated with a sharper decrease in cortisol levels over the course of the experiment, but given the small effect size and the absence of correction for multiple comparisons, this effect should be considered with great caution.

In summary, none of our lactate measures correlated significantly with any measure related to physical capacity, to fatigue, nor to any measure related to sleep, stress, or anxiety, emphasizing that dmPFC/dACC lactate influence relates to motivation and not to changes in capacity or other biological factors.

#### fMRI correlates of -SVch and DT

Remarkably, when looking at the neural correlates of negative SVch, there was no cluster surviving at a voxel-wise threshold of p < 0.05 family-wise error (FWE) corrected for multiple comparisons. However, when using a cluster-wise threshold of p < 0.05 FWE corrected for multiple comparisons, there was one single cluster centered in the dmPFC/dACC (Fig. S3-2A; x = -4, y = 26, z = 38; k = 266; p = 0.044). There was also one widespread network of activity correlating positively with DT at p < 0.001 FWE corrected for multiple comparisons at the voxel level, which included a big cluster in the dmPFC/dACC (Fig. S3-2B; x = -4, y = 24, z = 42; k = 1567; p < 0.001), a bilateral activation in the aIns, with the left aIns cluster extending in the left dorsolateral prefrontal cortex (dlPFC) which was also present bilaterally, possibly reflecting the involvement of the executive network with longer deliberation times [10, 18]. See also the full maps in [Neurovault](https://neurovault.org/collections/17029/) for more details.

#### Controlling For Possible Confounds Between Ech, SVch and DT Effects in the dmPFC/dACC

Crucially, the fact that dmPFC/dACC activity correlated with both Ech and DT and also tended to correlate negatively with SVch (Fig. 3C, Fig. S3-2A-B), even when these three parametric modulators of the choice period were split per task (Fig. S3-1), may be attributed to the rather big (20x20x20 mm^3^) dmPFC/dACC voxel that we used for ^1^H-MRS measurement in comparison with the clusters of activity (Fig. S1-2A; Fig. 3B). This voxel could overlap across several functional areas within the dmPFC/dACC, as has been suggested previously [19–21]. To assess this possibility, we extracted the cluster within the dmPFC/dACC corresponding to each of these regressors and looked at their overlap. Strikingly, there is a partial overlap between the three clusters (Fig. S3-2C-D). However, the DT cluster appears to be more dorsal (peak at x = -4, y = 24, z = 42), than the -SVch (x = -4, y = 26, z = 38) or the Ech (x = 0, y = 32, z = 32) clusters which overlapped considerably with each other (Fig. S3-2C), suggesting that they may be partially confounded at the behavioral and neural level. We therefore first verified whether variables were correlated at the behavioral level and found partial correlations between Ech and SVch (r = -0.081, p = 0.007, R² = 0.054) and between SVch and DT (r = -0.293, p < 0.001, R² =0.096), but Ech and DT were not significantly correlated to each other (r = -0.018, p = 0.325, R² = 0.015), suggesting that Ech was partially confounded with SVch at the behavioral level as well, although it was not confounded with DT. To clear out any potential confound for the neural correlates of our variables of interest, we therefore built two more GLM (GLM2 & GLM3) where we orthogonalized variables in the DT/SVch/Ech and in the Ech/SVch/DT order, respectively. GLM2 confirmed the significant correlation of both dmPFC/dACC and aIns activities with effort chosen (Fig. S3-3A; dmPFC/dACC: β = 0.397 ± 0.100, p < 0.001; aIns: β = 0.374 ± 0.087, p < 0.001), despite Ech being orthogonalized to both DT and SVch. Furthermore, this pattern also persisted for both physical (dmPFC/dACC: β = 0.420 ± 0.152; p = 0.007; aIns: β = 0.454 ± 0.120, p < 0.001) and mental (dmPFC/dACC: β = 0.374 ± 0.158; p = 0.021; aIns: β = 0.294 ± 0.140, p = 0.041) effort tasks. Moreover, in both GLM2 and GLM3 and despite orthogonalization, both dmPFC/dACC and aIns correlated negatively to SVch (Fig. S3-3A-B) and positively to DT (Fig. S3-3A-B), confirming that the dmPFC/dACC (at least in the location where we measured ^1^H-MRS) correlates with all three variables.

Since we observed that dmPFC/dACC and aIns activities also correlated with SVch and DT, we evaluated whether the correlation between neural activity and behavior were exclusively associated with the individual parameter estimate corresponding to the Ech regressor as shown in Fig. 4 or whether SVch and/or DT regression estimates in the dmPFC/dACC or in the aIns would also correlate with inter-individual differences in HE choices proportion. Our findings revealed no significant correlation between the regression estimate for SVch or DT and the overall proportion of HE choices (Fig. S4-1C-D), neither in dmPFC/dACC (SVch: r = -0.098, p = 0.452; DT: r = -0.012, p = 0.929) nor in aIns (SVch: r = -0.01, p = 0.938; DT: r = -0.012, p = 0.930). The lack of significant correlation also held when looking specifically at HPE choices (Fig. S4-1C-D) and dmPFC/dACC activity for SVch (r = -0.232, p = 0.070) and for DT (r = -0.026, p = 0.839) and for aIns activity for SVch (r = -0.101, p = 0.441) and DT (r = -0.043, p = 0.743). The same was also true when looking at HME choices, and kEp and kEm parameters (Fig. S4-1C-D). This lack of correlation underscores the specificity of the neural response to the effort component for explaining differences in motivated behavior, rather than to other aspects of decision-making which also drive dmPFC/dACC and aIns activities.

#### Effort Chosen or Effort Cost of the High Effort Option?

Lastly, to control that the dmPFC/dACC was driving high-effort choices, rather than computing the cost of the high-effort option, we designed a fourth GLM (GLM4) where low- and high-effort choices were modeled as separate regressors, each parametrically modulated by the level of the high-effort option. Therefore, we confirmed that both dmPFC/dACC and aIns activities increased with the high effort level only when the high effort was picked (Fig. S3-4). This result confirms that both dmPFC/dACC and aIns activities were not correlated with the effort cost of the high effort (HE) option, but with the difficulty of the effort chosen, which further confirms that dmPFC/dACC and aIns activities are predictive of the tendency to select the HE option within a given individual.

### Supplementary Discussion

Interestingly, on top of its correlation with the level of the chosen effort, as expected from the literature, the dmPFC/dACC also correlated positively with deliberation times [10, 18, 22, 23] and negatively with subjective value [24–26] in our own data. However, the encoding of deliberation times (representing decision costs) in the dmPFC/dACC was slightly more dorsal than the correlates of effort chosen and the subjective value of the chosen option (decision value) which were more ventral, in agreement with a ventro-dorsal gradient between value and conflict encoding in this region [21]. It is therefore rather possible that different subparts of the dmPFC/dACC relate to different functions [19]. It would be interesting that future studies explore whether lactate levels vary across the different parts of the dmPFC/dACC and whether these changes relate to a differential impact on motivated behavior depending on their location.

Prospect theory because prospect theory states 1) that individuals display loss aversion and weigh losses higher than equivalent monetary gains and 2) that individuals are more risk seeking in the loss domain than in the gain domain [27]. These two proposals imply that, in our task, individuals should pick the HE option more often in the loss than in the gain domain because the same monetary amounts should have more weight in the loss than in the gain domain and because the probability of failure is slightly higher for high than for low efforts due to difficulty, although we reduced the risk of failure as much as possible in our task. Instead, our results suggest that individuals weigh gains more than losses as they picked up the HE option more often in the gain than in the loss domains and were more sensitive to gains than to losses. In agreement with a vast and rising literature contesting the loss aversion bias [28–30], our results thereby question some of prospect theory’s predictions regarding loss aversion and risk seeking in the loss domain. Interestingly, several studies suggest that for “low” amounts (typically < 50€/CHF/$/etc. as in our experiment), individuals value gains more than losses [28, 30–32]. Loss aversion, the fact of weighing equivalent losses more than equivalent gains, would emerge only for “high” amounts (typically > 1000$) as the ones used in the original experiments of the tenants of the prospect theory [28], *i.e.* when the loss is really consequential for one’s life and survival. Our results, which show faster choices and more HE choices in the gain than in the loss domain, are consistent with opponency theories of approach/avoidance where the expectation of more positive stimuli tends to trigger faster and more intense approach answers due to dopamine release, while the expectation of more negative stimuli tends to trigger slower reactions based on inhibition and avoidance, potentially triggered by serotonin release [8, 12].

### Supplementary Tables

| **Female/Male (N)** | **Age (years)** | **Weight (kg)** | **Height (m)** | **BMI** | **Level of studies** |
| --- | --- | --- | --- | --- | --- |
| 36/27 | 29.7 ± 0.49 | 66.4 ± 1.37 | 1.71±0.03 | 22.54±0.50 | 6.27±0.21 |

#### Table S1: Demographic characteristics of the (included) participants (n = 63).

The level of studies is indicated based on the International Standard Classification of Education (ISCED) which is a range going from 0 to 8. Results are mean ± SEM.

### Supplementary Figures

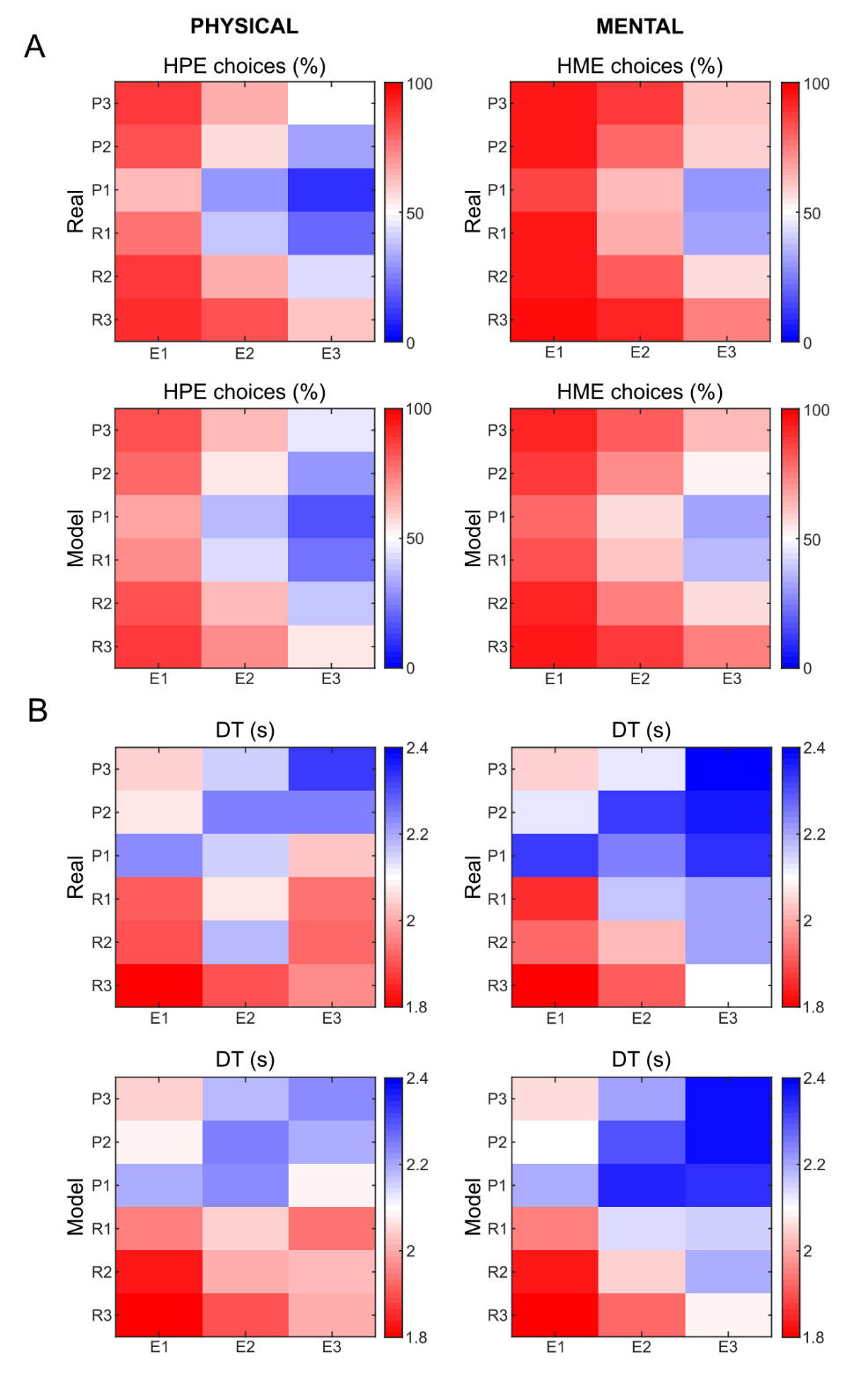

#### Figure S1-1: Behavioral results.

**A]** Proportion of high physical effort (HPE, left panels) and high mental effort (HME, right panels) choices selected by the subjects (top panels) or predicted by our computational model (bottom panels) in function of the monetary incentives expressed as rewards (R1/R2/R3) or punishments (P1/P2/P3) and of the effort level (E1/E2/E3) of the varying option. **B]** Deliberation times (DT) (top panels) and fitted DT (bottom panels) in seconds (s) in function of the monetary incentives and of the effort level of the varying option. Note that in all graphs a higher level of incentive (reward R or punishment P) means that it was more valuable (higher difference with the fixed option) while a higher level of effort means that the effort was more difficult.

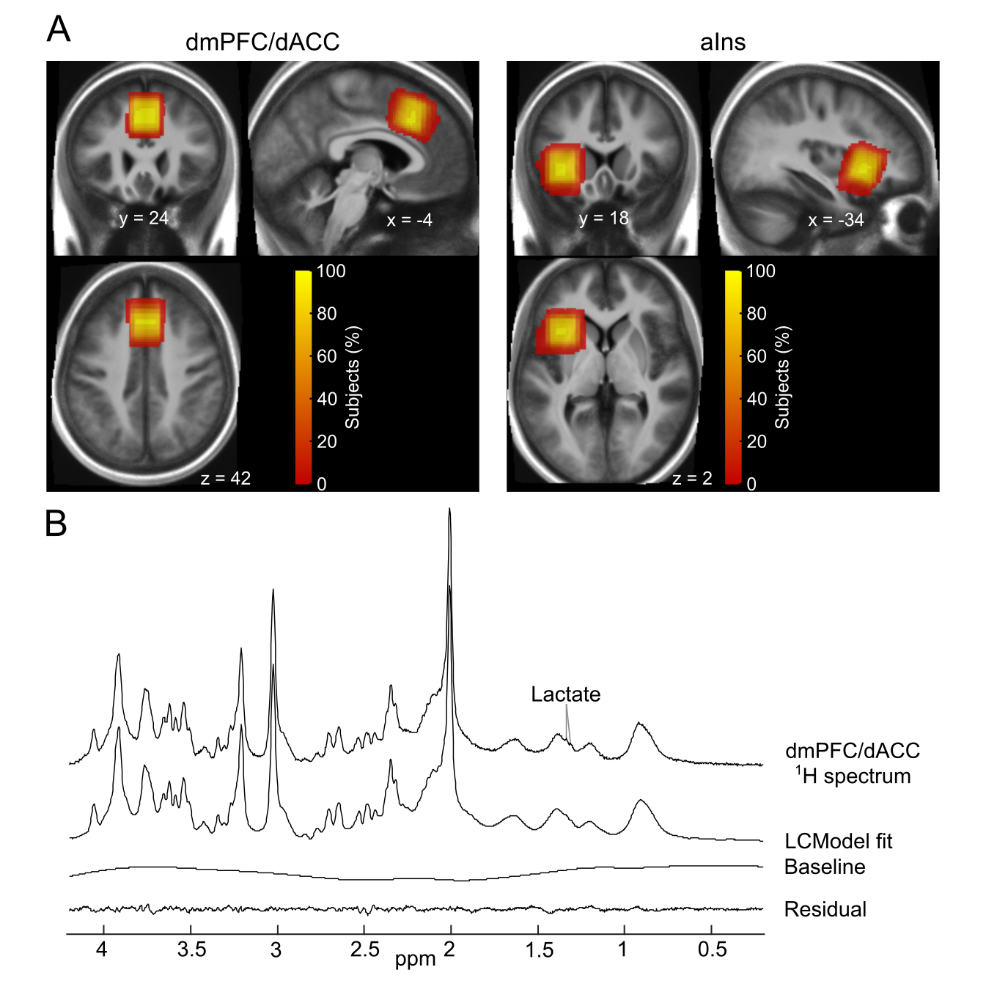

#### Figure S1-2: Density map of the ^1^H-MRS voxels.

**A**] Proton magnetic resonance spectroscopy (^1^H-MRS) density maps representing the proportion of subjects where each voxel is present across individuals overlaid on the average anatomy of the subjects included in the study (N = 63). All the data is represented in MNI space. Note that the aIns map contains lesser subjects (N = 60) than the dmPFC/dACC map (N = 63) because the aIns MRS measurement could not be acquired in some subjects. **B**] Representative ^1^H-MRS spectrum acquired with the semi-adiabatic SPECIAL sequence at 7 Tesla, as well as the corresponding LCModel spectral fit, baseline and residual fit. The arrow highlights the lactate doublet.

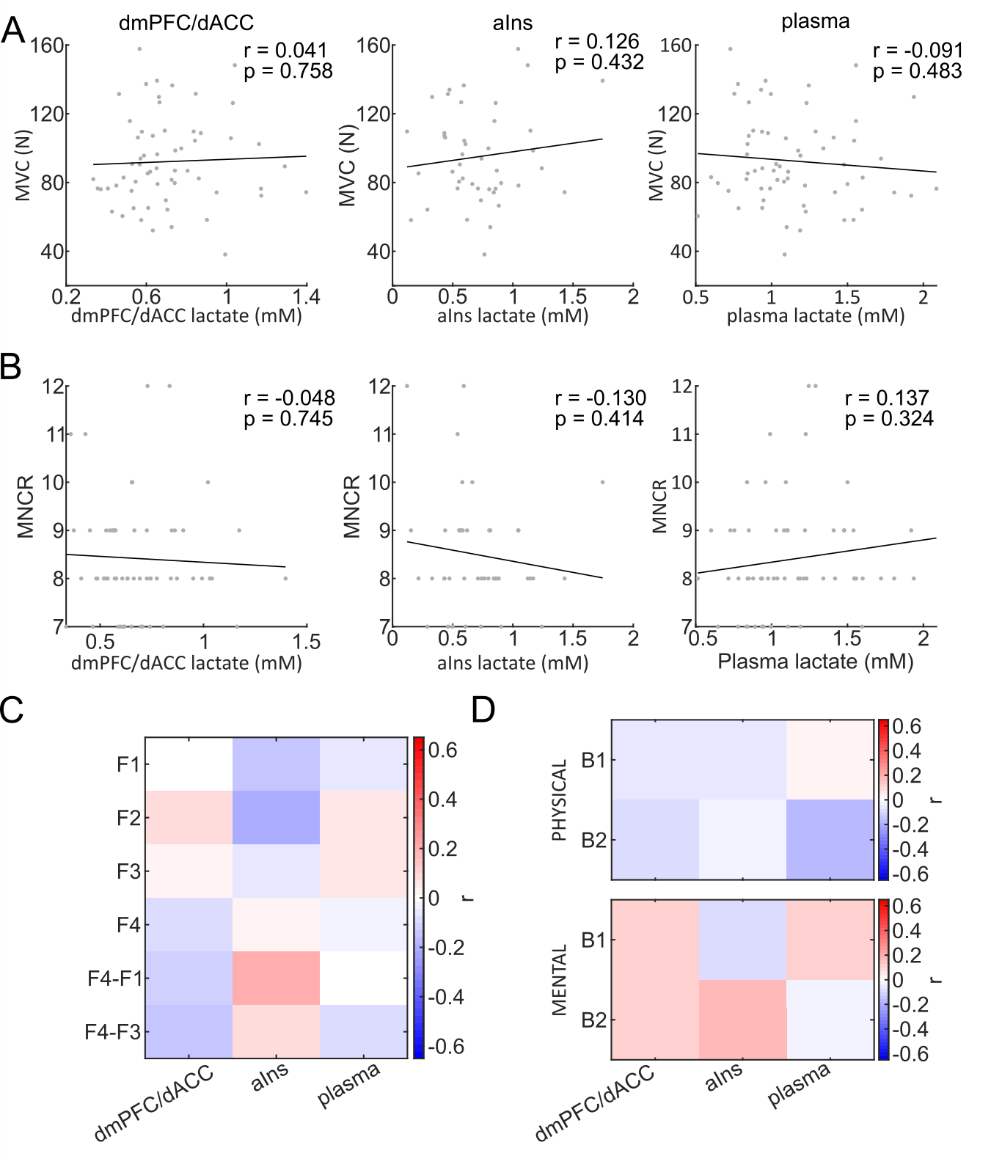

#### Figure S2-1: Correlations between plasma and brain lactate levels and MVC, subjective fatigue and maximal performance.

**A**] Maximum voluntary contraction (MVC) force produced in Newtons (N) during the initial calibration in function of individual levels of lactate in the dmPFC/dACC (left), the aIns (middle), or the plasma (right). **B**] Maximum number of correct responses (MNCR) produced during the initial calibration in function of individual levels of lactate in the dmPFC/dACC (left), the aIns (middle), or the plasma (right). **C**] Heatmap displaying Pearson correlation coefficients between subjective fatigue ratings before (F1) or after (F2) the ^1^H-MRS acquisition, before (F3) or after the fMRI task (F4), and difference between F4 and F1 or between F4 and F3 and dmPFC/dACC, aIns, or plasma lactate levels. None of the correlations is significant. **D**] Difference in maximal performance after versus before each of the two blocks (B) of each task in function of lactate in the plasma, the dmPFC/dACC, or the aIns. None of the correlations is significant.

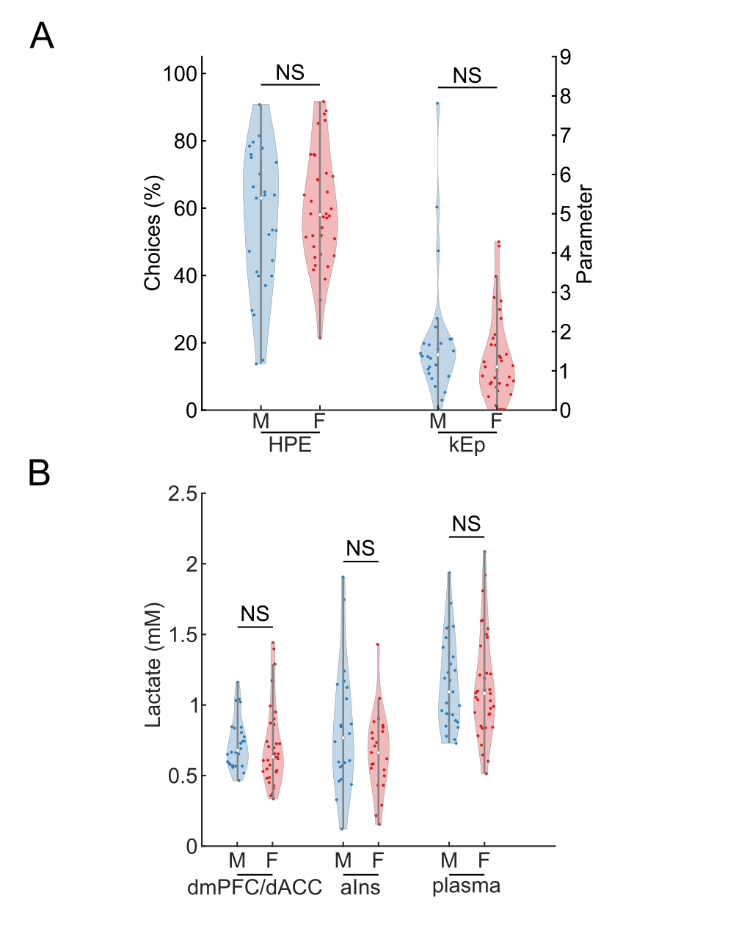

#### Figure S2-2: Absence of sex differences for lactate concentrations and motivation-related measures.

**A**] No significant (NS) difference between males (M) and females (F) for the proportion of high physical effort (HPE) choices or for the kEp parameter. **B**] No significant (NS) differences between males (M) and females (F) for dmPFC/dACC, aIns and plasma lactate levels.

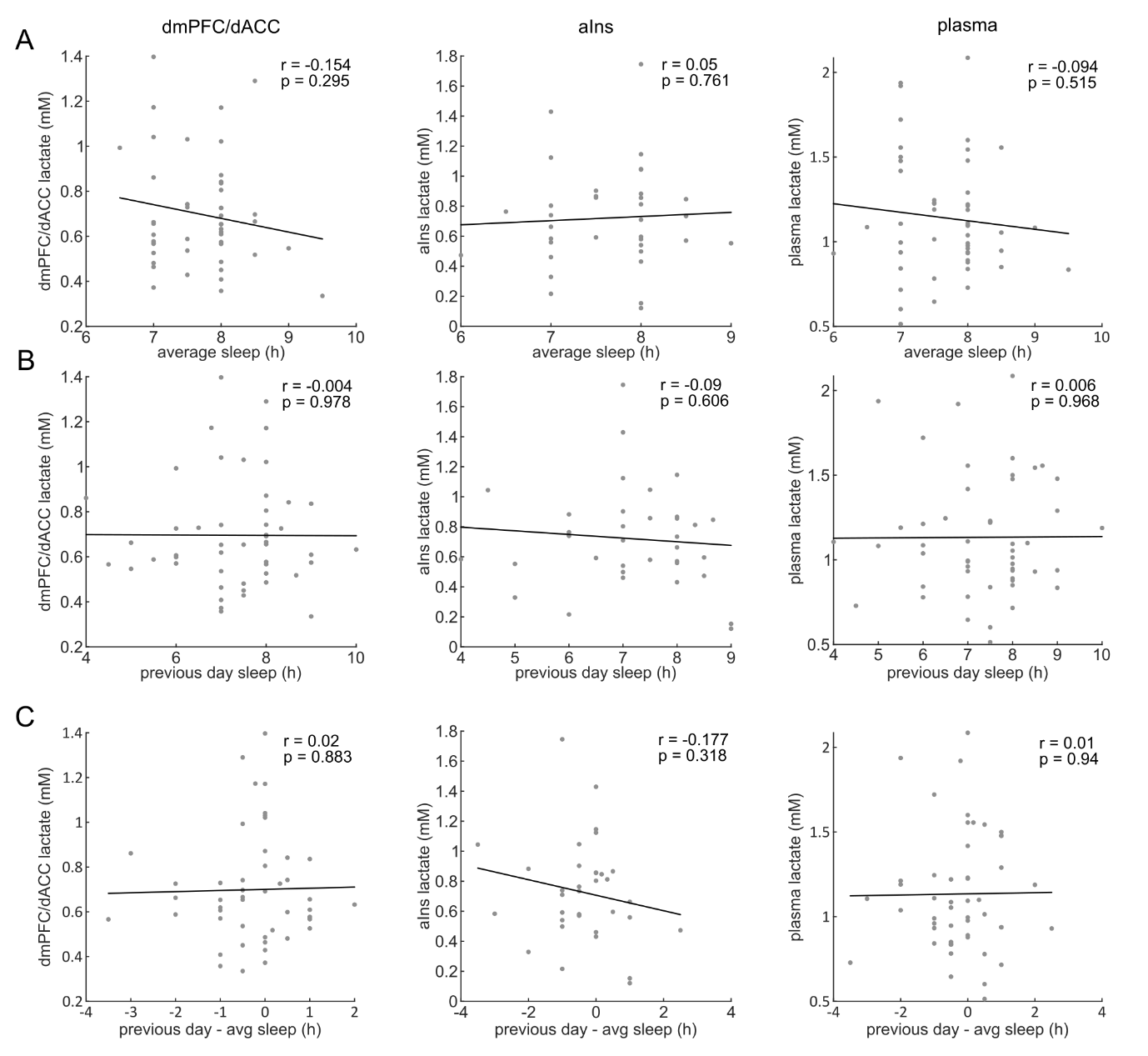

#### Figure S2-3: Correlations between plasma and brain lactate levels and sleep duration.

Plasma, dmPFC/dACC and aIns lactate levels in function of average sleep duration (**A**), sleep duration the day preceding the experiment (**B**) and the difference between the sleep the day preceding the experiment and the average duration of sleep (**C**).

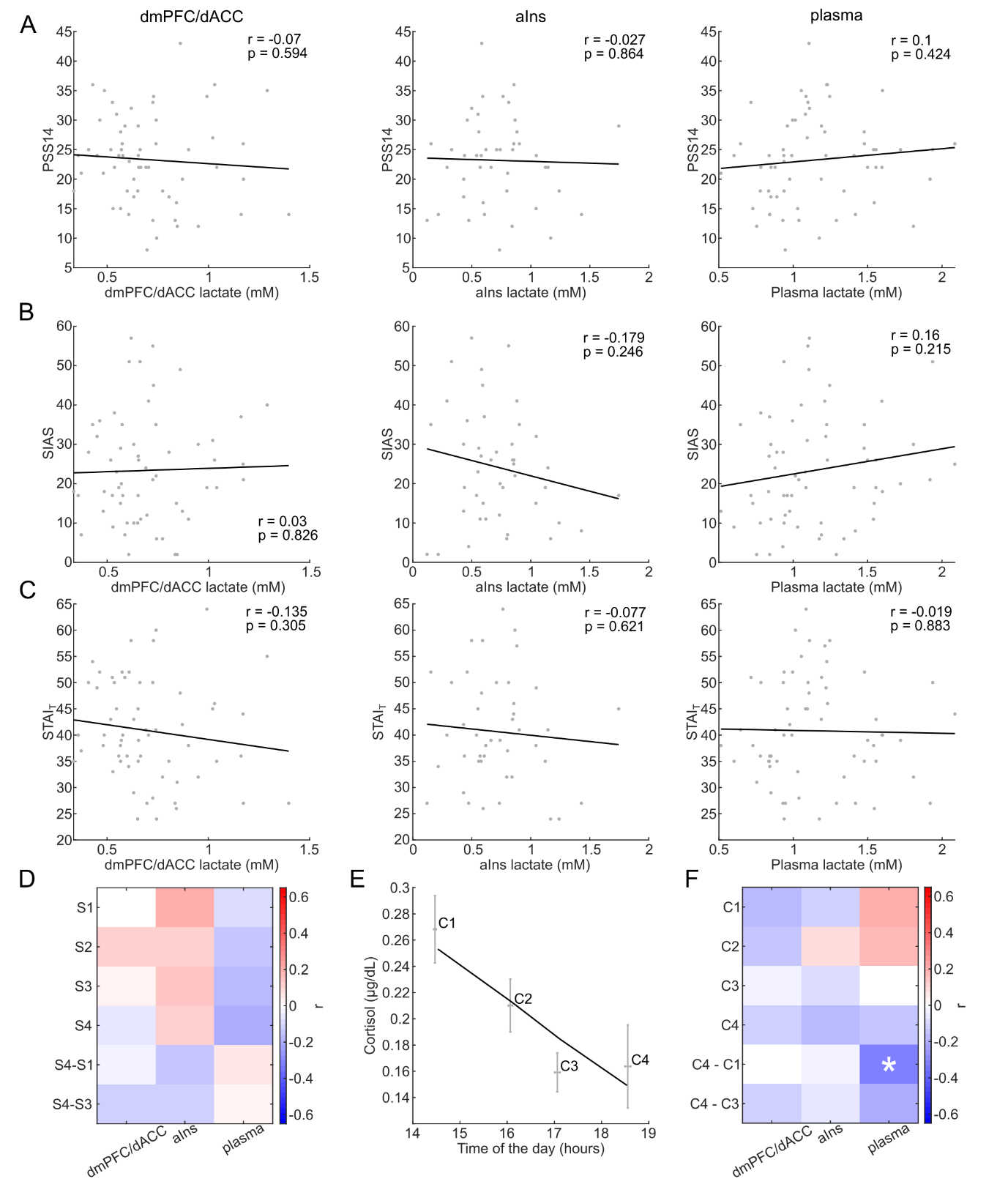

#### Figure S2-4: Correlations between plasma and brain lactate levels and stress-related measures.

**A**] Perceived Stress Scale (PSS-14) score in function of dmPFC/dACC (left), aIns (middle) or plasma (right) levels. **B**] Social Interaction Anxiety Scale (SIAS) score in function of dmPFC/dACC (left), aIns (middle) or plasma (right) levels. **C**] Spielberger’s State-Trait Anxiety Inventory – trait (STAI_T_) in function of dmPFC/dACC (left), aIns (middle) or plasma (right) levels. **D**] Stress ratings before (S1) or after (S2) the ^1^H-MRS acquisition, before (S3) or after the fMRI task (S4), and difference between S4 and S1 or between S4 and S3 and dmPFC/dACC, aIns, or plasma lactate levels. None of the correlation is significant. **E**] Salivary cortisol across the experiment. Cortisol measures were taken before (C1) and after (C2) the ^1^H-MRS acquisition, and before (C3) and after (C4) the MRI acquisition. **F**] Salivary cortisol across the duration of the experiment (C1-4), and difference between C4 and C1 or between C4 and C3 in function of dmPFC/dACC, aIns, or plasma lactate levels. The only significant correlation was between C4-C1 and lactate plasma levels (r = -0.311; p = 0.019 uncorrected for multiple comparisons).

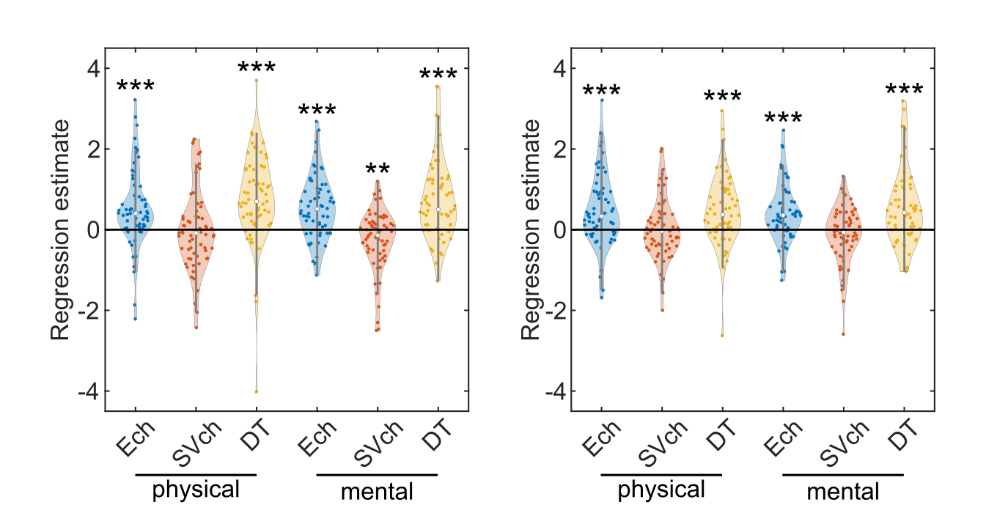

#### Figure S3-1: dmPFC/dACC and aIns activity correlation with Ech, SVch and DT separately for each task.

Regression estimates for effort chosen (Ech), the subjective value of the chosen option (SVch) and deliberation time (DT) during the choice period within the ^1^H-MRS dmPFC/dACC (N = 63, left panel) and the ^1^H-MRS aIns (N = 60, right panel) for each individual (displayed in circles) looking independently for each task. The dmPFC/dACC activity is positively associated to Ech in both the physical (β = 0.516 ± 0.123, p < 0.001) and the mental (β = 0.551 ± 0.098, p < 0.001) effort tasks, as well as with DT in both the physical (β = 0.764 ± 0.141, p < 0.001) and the mental (β = 0.712 ± 0.123, p < 0.001) effort tasks. It also correlated negatively with SVch in the mental (β = -0.283 ± 0.098, p = 0.005), but not in the physical (β = -0.012 ± 0.126, p = 0.927) task. The aIns correlated positively with Ech in both physical (β = 0.471 ± 0.117, p < 0.001) and mental (β = 0.411 ± 0.094, p < 0.001) effort tasks, as well as with DT in both physical (β = 0.466 ± 0.112, p < 0.001) and mental effort tasks (β = 0.534 ± 0.120, p < 0.001), but it did not significantly correlate to SVch, neither in the physical (β = 0.025 ± 0.100, p = 0.806), nor in the mental (β = -0.128 ± 0.091, p = 0.165) effort task. Significance of Student t-tests against zero: ***p<0.001; **p<0.01; *p<0.05.

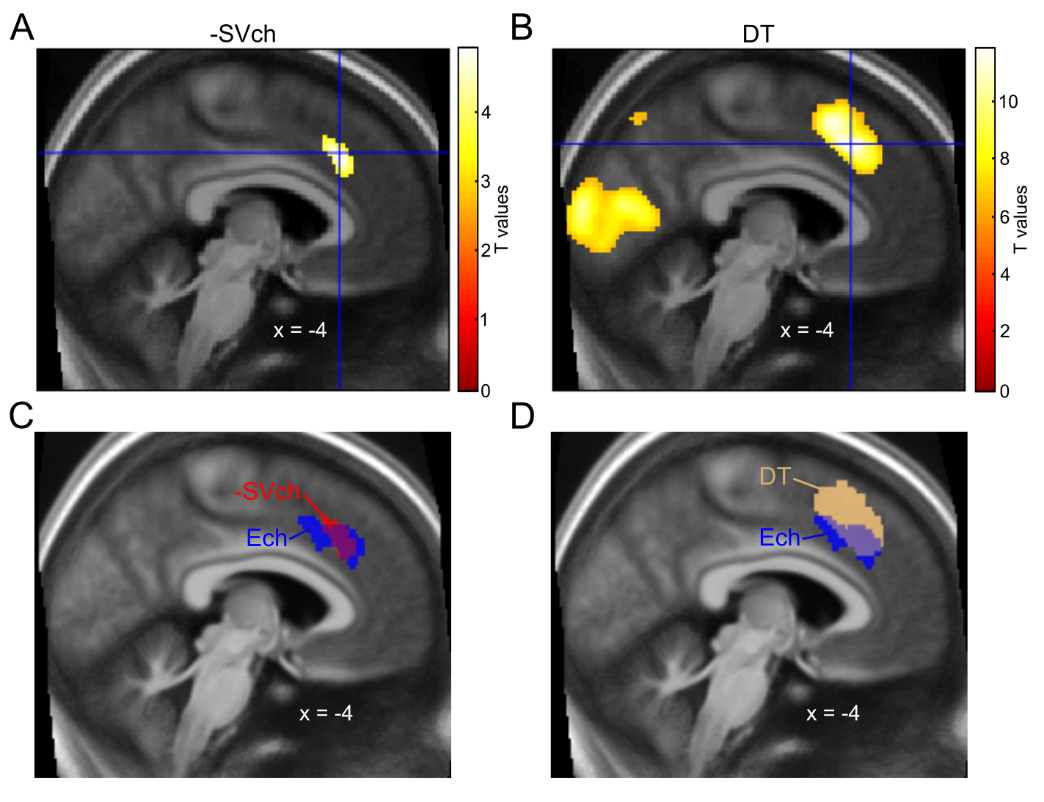

#### Figure S3-2: Neural correlates of -SVch and DT and overlap between Ech, -SVch and DT dmPFC/dACC clusters.

**A**] Negative neural correlates of the subjective value of the chosen option (SVch) across tasks. A t.test against zero across participants (N = 63) reveals only one cluster, centered in the dmPFC/dACC (x = -4, y = 26, z = 38; k = 266; p = 0.044), voxel-wise thresholded at p < 0.001 uncorrected for multiple comparisons but cluster-wise thresholded at p < 0.05 after family-wise error (FWE) correction for multiple comparisons. Note that no cluster survives with a voxel-wise threshold of p < 0.05 after FWE correction for multiple comparisons. **B**] Neural correlates of deliberation times (DT) across tasks. A t.test against zero across participants (N = 63), voxel-wise thresholded at p < 0.001 after FWE correction for multiple comparisons, reveals a widespread network including important clusters in the dmPFC/dACC (x = -4, y = 24, z = 42; k = 1567; p < 0.001) highlighted in the figure, as well as in the left (x = -48, y = 22, z = 28; k = 2543; p < 0.001) and right (x = 42, y = 20, z = 42; k = 319; p < 0.001) dorsolateral prefrontal cortex (dlPFC) and in the left (x = -34, y = 18, z = 0; k = 2543; p < 0.001) and right (x = 34, y = 24, z = -2; k = 345; p < 0.001) aIns. **C**] Overlap between the effort chosen (Ech, in blue) and the negative subjective value (-SVch, in red) dmPFC/dACC clusters. **D**] Overlap between the effort chosen (Ech, in blue) and the deliberation times (DT, in beige) dmPFC/dACC clusters. All fMRI clusters are overlaid on the anatomical average of the 63 subjects included in the study in MNI space.

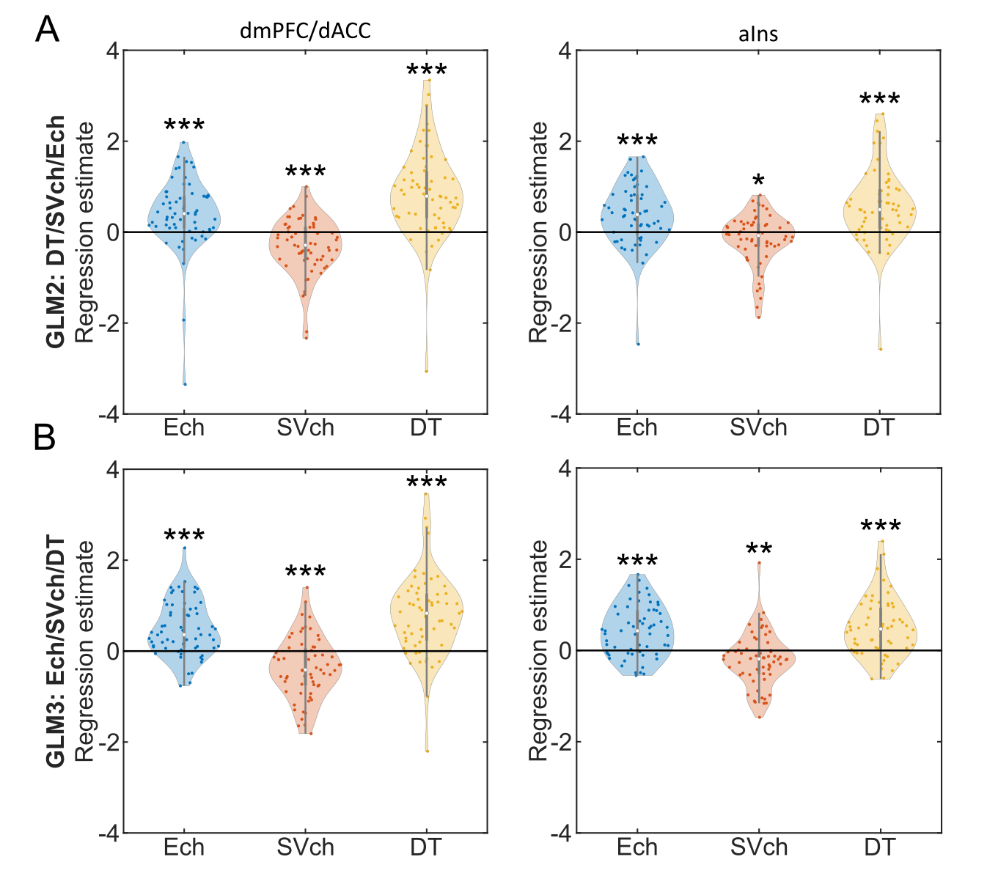

#### Figure S3-3: dmPFC/dACC (left panel) and aIns (right panel) activity regression estimates for effort chosen (Ech), subjective value of the chosen option (SVch) and deliberation time (DT) when variables are orthogonalized.

Data has been extracted in the dmPFC/dACC (left column) and in the aIns (right column) for [**A**] GLM2 where variables are orthogonalized in the DT/SVch/Ech order and [**B**] for GLM3 where variables are orthogonalized in the Ech/SVch/DT order. In all panels, the violin plots represent the regression estimates for each regressor included in the GLM at the time of choice (N = 63). Abbreviations mean level of the effort chosen (Ech), subjective value of the chosen option (SVch) and deliberation time (DT). Significance levels of t-test against zero: ***p<0.001; **p<0.01; *p<0.05.

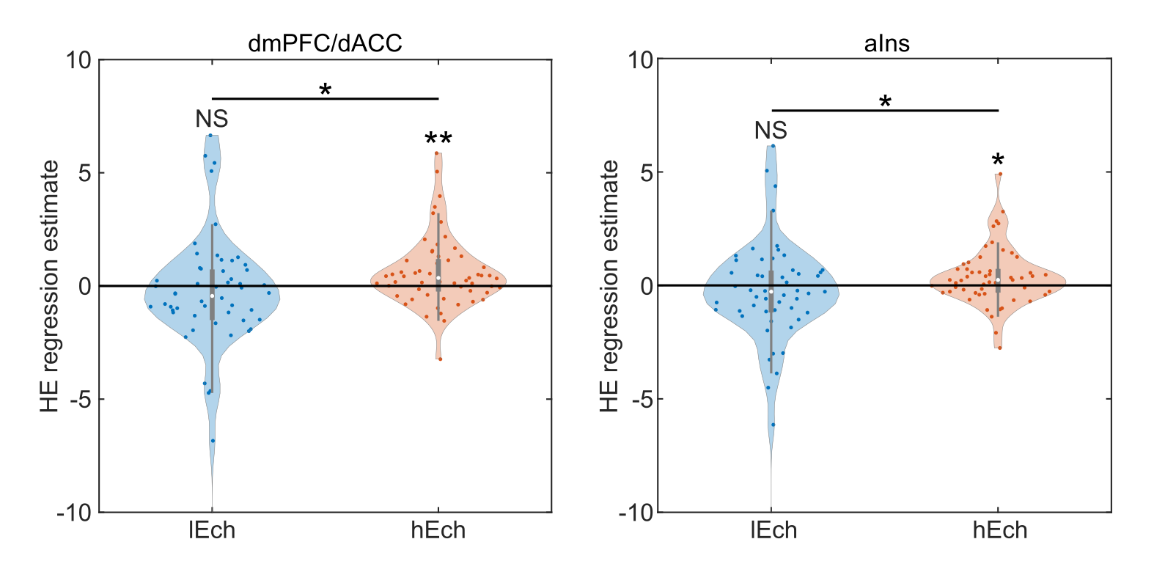

#### Figure S3-4: Neural activity in function of effort difficulty and choice.

The dmPFC/dACC (left panel) and aIns (right panel) regression estimate for high effort (HE) level was estimated independently for the trials when the low effort was chosen (lEch) and the trials when the high effort was chosen (hEch). The two regression estimates were compared with a paired t-test. Significance levels: ***p<0.001, **p<0.01, *p < 0.05, NS for not significant. Statistics and figures display the results after removing outliers defined as those who have a regression estimate further than the median ± 3*SD.

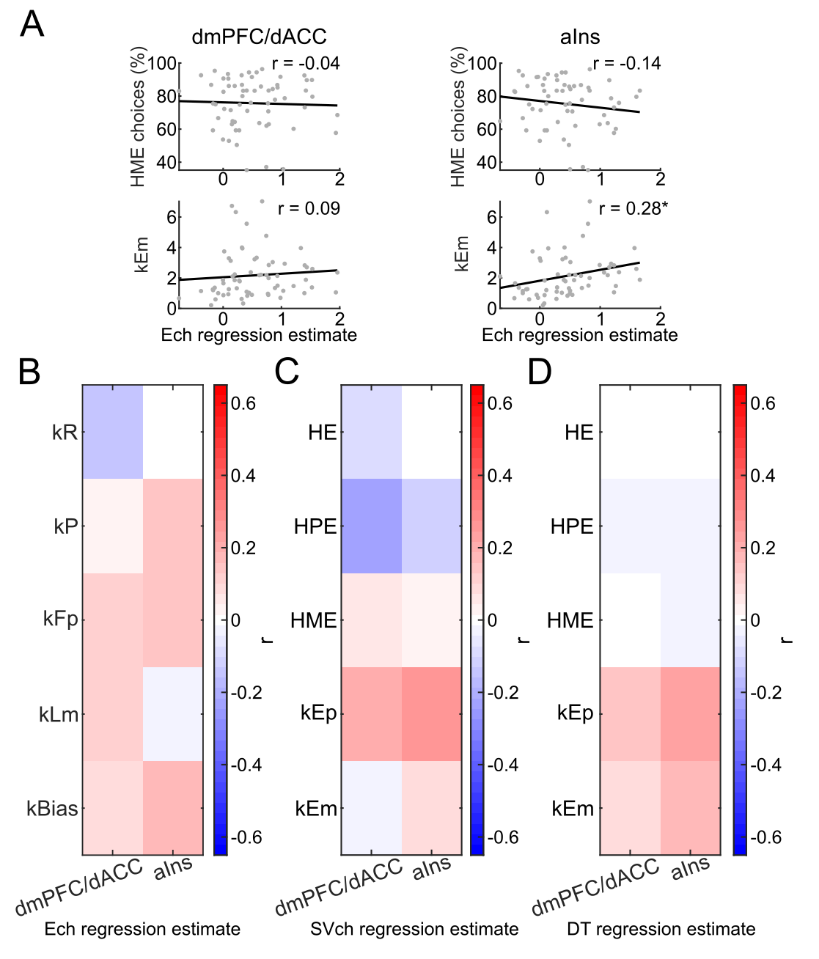

#### Figure S4-1: Correlation between dmPFC/dACC and aIns activity regression estimates during choice and behavioral motivation measures.

**A**] Scatter plots depicting the correlation between dmPFC/dACC (left panels) and aIns (right panels) effort chosen (Ech) regression estimates and the proportion of high mental effort (HME) choices (upper panels) and the sensitivity to mental effort kEm (bottom panels). Each point represents an individual participant. Significance levels: ***p<0.001; **p<0.01; *p<0.05. **B**] Heatmap depicting Pearson correlation coefficients between model parameters (kR, kP, kFp, kLm and kBias) against dmPFC/dACC and aIns effort chosen (Ech) regression estimate during the choice period. None of the correlations is significant. **C-D**] Heatmap representing Pearson correlation coefficients between the proportion of high effort choices globally (HE), in the physical effort task (HPE) and in the mental effort task (HME) as well as effort sensitivities for physical effort (kEp) and mental effort (kEm) against dmPFC/dACC and aIns regression estimates for [**C**] the subjective value of the chosen option (SVch) and [**D**] for deliberation times (DT). None of the correlations displayed is significant.

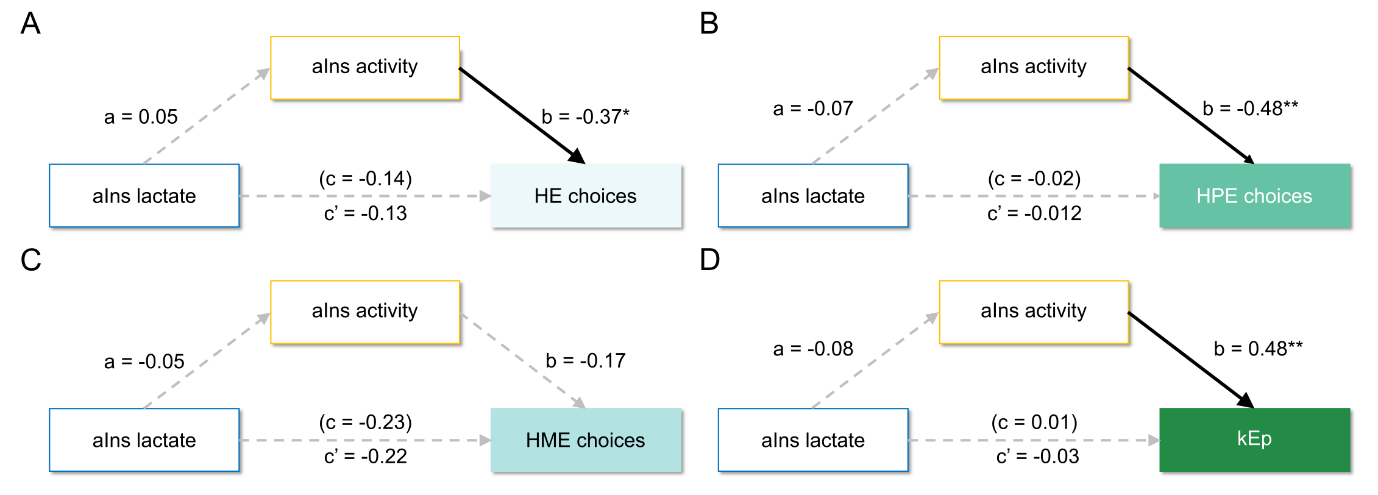

#### Figure S5-1: Mediation analysis of aIns lactate on motivated behavior, via aIns activity.

Significant paths are depicted with continuous black lines, while non-significant paths are represented with dotted grey lines. Path c represents the coefficient for the direct path before considering the mediator, while path c’ represents the coefficient for the direct path after considering the mediator in the same equation, and path a and b are the coefficients for the indirect pathway (*i.e.* the mediation). Significance levels for each path: ***p<0.001; **p<0.01; *p<0.05. **A**] Total high-effort (HE) choices: Path analysis testing if aIns lactate levels influence HE choices mediated by aIns fMRI regression estimate for effort chosen (N = 43). **B**] High physical effort (HPE) choices: Path analysis testing if aIns lactate levels influence HPE choices mediated by aIns fMRI regression estimate for effort chosen (N = 44). **C**] High mental effort (HME) choices: Path analysis testing if aIns lactate levels influence HME choices mediated by aIns fMRI regression estimate for effort chosen (N = 43). **D**] Sensitivity to physical effort kEp: Path analysis testing if aIns lactate levels influence kEp, mediated by the aIns fMRI regression estimate for effort chosen (N = 42).

### Supplementary References

1.	Lesage F-X, Berjot S, Deschamps F. Psychometric properties of the French versions of the Perceived Stress Scale. Int J Occup Med Environ Health. 2012;25:178–184.

2.	Spielberger CD, Bruchon-Schweitzer M, Paulhan I. STAI-Y: Inventaire d’anxiété état-trait forme Y. Paris, France: Éditions du centre de psychologie appliquée, DL 1993; 1993.

3.	Mattick RP, Clarke JC. Social Interaction Anxiety Scale. 2012.

4.	Westbrook A, Lamichhane B, Braver T. The Subjective Value of Cognitive Effort is Encoded by a Domain-General Valuation Network. The Journal of Neuroscience. 2019:3071–18.

5.	Westbrook A, Kester D, Braver TS. What Is the Subjective Cost of Cognitive Effort? Load, Trait, and Aging Effects Revealed by Economic Preference. PLoS ONE. 2013;8:e68210.

6.	Daunizeau J, Adam V, Rigoux L. VBA: A Probabilistic Treatment of Nonlinear Models for Neurobiological and Behavioural Data. PLoS Comput Biol. 2014;10:e1003441.

7.	Wilson RC, Collins AG. Ten simple rules for the computational modeling of behavioral data. eLife. 2019;8:e49547.

8.	Shadmehr R, Reppert TR, Summerside EM, Yoon T, Ahmed AA. Movement Vigor as a Reflection of Subjective Economic Utility. Trends in Neurosciences. 2019;42:323–336.

9.	Lebreton M, Abitbol R, Daunizeau J, Pessiglione M. Automatic integration of confidence in the brain valuation signal. Nat Neurosci. 2015;18:1159–1167.

10.	Clairis N, Pessiglione M. Value, confidence, deliberation: a functional partition of the medial prefrontal cortex demonstrated across rating and choice tasks. J Neurosci. 2022;42:5580–5592.

11.	De Martino B, Fleming SM, Garrett N, Dolan RJ. Confidence in value-based choice. Nat Neurosci. 2013;16:105–110.

12.	Boureau Y-L, Dayan P. Opponency revisited: competition and cooperation between dopamine and serotonin. Neuropsychopharmacology. 2011;36:74–97.

13.	Steverson K, Chung H-K, Zimmermann J, Louie K, Glimcher P. Sensitivity of reaction time to the magnitude of rewards reveals the cost-structure of time. Sci Rep. 2019;9:20053.

14.	Chow LS, Gerszten RE, Taylor JM, Pedersen BK, van Praag H, Trappe S, et al. Exerkines in health, resilience and disease. Nat Rev Endocrinol. 2022;18:273–289.

15.	Hladky SB, Barrand MA. Metabolite Clearance During Wakefulness and Sleep. Handb Exp Pharmacol. 2019;253:385–423.

16.	Lundgaard I, Lu ML, Yang E, Peng W, Mestre H, Hitomi E, et al. Glymphatic clearance controls state-dependent changes in brain lactate concentration. J Cereb Blood Flow Metab. 2017;37:2112–2124.

17.	Chamaa F, Magistretti PJ, Fiumelli H. Astrocyte-derived lactate in stress disorders. Neurobiol Dis. 2024;192:106417.

18.	Clairis N, Pessiglione M. Value Estimation versus Effort Mobilization: A General Dissociation between Ventromedial and Dorsomedial Prefrontal Cortex. J Neurosci. 2024;44.

19.	Clairis N, Lopez-Persem A. Debates on the dorsomedial prefrontal/dorsal anterior cingulate cortex: insights for future research. Brain. 2023:awad263.

20.	Jahn A, Nee DE, Alexander WH, Brown JW. Distinct Regions within Medial Prefrontal Cortex Process Pain and Cognition. J Neurosci. 2016;36:12385–12392.

21.	Kolling N, Wittmann MK, Behrens TEJ, Boorman ED, Mars RB, Rushworth MFS. Value, search, persistence and model updating in anterior cingulate cortex. Nature Neuroscience. 2016;19:1280–1285.

22.	Grinband J, Savitskaya J, Wager TD, Teichert T, Ferrera VP, Hirsch J. The dorsal medial frontal cortex is sensitive to time on task, not response conflict or error likelihood. Neuroimage. 2011;57:303–311.

23.	McGuire JT, Botvinick MM. Prefrontal cortex, cognitive control, and the registration of decision costs. Proc Natl Acad Sci U S A. 2010;107:7922–7926.

24.	Bartra O, McGuire JT, Kable JW. The valuation system: A coordinate-based meta-analysis of BOLD fMRI experiments examining neural correlates of subjective value. NeuroImage. 2013;76:412–427.

25.	Le Bouc R, Pessiglione M. A neuro-computational account of procrastination behavior. Nat Commun. 2022;13:5639.

26.	Yao Y-W, Song K-R, Schuck NW, Li X, Fang X-Y, Zhang J-T, et al. The dorsomedial prefrontal cortex represents subjective value across effort-based and risky decision-making. Neuroimage. 2023;279:120326.

27.	Kahneman D, Tversky A. Choices, values, and frames. American Psychologist. 1984;39:341–350.

28.	Yechiam E. Acceptable losses: the debatable origins of loss aversion. Psychol Res. 2019;83:1327–1339.

29.	Gal D, Rucker DD. The Loss of Loss Aversion: Will It Loom Larger Than Its Gain? Journal of Consumer Psychology. 2018;28:497–516.

30.	Mukherjee S, Sahay A, Pammi VSC, Srinivasan N. Is loss-aversion magnitude-dependent? Measuring prospective aﬀective judgments regarding gains and losses. Judgment and Decision Making. 2017;12:81–89.

31.	Harinck F, Van Dijk E, Van Beest I, Mersmann P. When Gains Loom Larger Than Losses: Reversed Loss Aversion for Small Amounts of Money. Psychol Sci. 2007;18:1099–1105.

32.	Ert E, Erev I. On the Descriptive Value of Loss Aversion in Decisions under Risk: Six Clarifications. Rochester, NY: Social Science Research Network; 2007.
